# Distinct aging-vulnerable trajectories of motor circuit functions in oxidation- and temperature-stressed *Drosophila*

**DOI:** 10.1101/2020.08.19.257832

**Authors:** Atulya Srisudarshan Ram Iyengar, Hongyu Ruan, Chun-Fang Wu

## Abstract

We examined several sensory-motor processing circuits in *Drosophila* across the lifespan and uncovered distinctive age-resilient and age-vulnerable trajectories in their established functional properties. We observed relatively little deterioration toward the end of lifespan in the giant-fiber (GF) and downstream circuit elements responsible for the jump-and-flight escape reflex. In contrast, we found substantial age-dependent modifications in the performance of GF inputs and other circuits driving flight motoneuron activities. Importantly, in high temperature (HT)-reared flies (29 °C), the characteristic age-dependent progression of these properties was largely maintained, albeit over a compressed time scale, lending support for the common practice of expediting *Drosophila* aging studies by HT rearing. We discovered shortened lifespans in *Cu*^*2+*^*/Zn*^*2+*^ *Superoxide Dismutase 1* (*Sod*) mutant flies were accompanied by alterations distinct from HT-reared flies, highlighting differential effects of oxidative vs temperature stressors. This work also establishes several age-vulnerable parameters that may serve as quantitative neurophysiological landmarks for aging in *Drosophila*.

## Introduction

Aging nervous systems manifest a progressive functional decline. Invertebrate preparations offer experimentally tractable, simpler nervous systems with individually identifiable neurons to reveal the molecular and cellular bases of age-related neurophysiological changes (***Janse et al., 1986***; ***Atwood, 1992; Kenyon, 2001; Yeoman and Faragher, 2001; Pletcher et al., 2002; Toivonen and Partridge, 2009***). In *Drosophila*, characteristic declines over the lifespan have been documented in motor coordination (***Leffelaar and Grigliatti, 1983; Gargano et al., 2005***) and in higher functions, such as learning and memory (***Tamura et al., 2003; Yamazaki et al., 2007***). However, fewer studies have directly assessed cellular physiological decline underlying changes in behavioral performance across the lifespan (e.g. ***Martinez et al. ((2007***)). It is therefore important to identify the neural circuits and associated behavioral outputs that are prone to age-related modifications and determine whether specific neuronal elements are differentially vulnerable in this process. Given that environmental and genetic contributions must be both considered in studies of the aging process (***Miquel et al., 1976; Curtsinger et al., 1995; Mair et al., 2003***), it is equally important to examine how age-progression of circuit function is differentially modified by extrinsic environmental and intrinsic genetic factors. Such analyses may also uncover salient physiological parameters suitable for indexing aging progression.

This study examines several motor-pattern circuits in *Drosophila* whose outputs converge on an identified, well-described motor neuron driving the activity of a subset of flight muscles (dorsal longitudinal muscles, DLM a-e, see ***Levine (1973))***. Thus the DLM readout, driven by different motor circuits, enabled us to determine how age-dependent changes in different categories of motor performance are modified by either high-temperature rearing (29 °C, leading to a shortened lifespan), or oxidative stress in *Cu*^*2+*^*/Zn*^*2+*^ *Superoxide Dismutase 1* (*Sod*) mutants (defective in a major free radical-scavenging enzyme, see ***Campbell et al. ((1986)***; ***Phillips et al. ((1989))***. Rearing flies at 29 °C is a widely adopted protocol for experimental expedience in *Drosophila* aging and neurodegeneration studies (e.g. ***Min and Benzer (1997)***; ***Kang et al. ((2002)***; ***Simon et al. ((2003)***; ***Biteau et al. (2010)***; ***Sudmeier et al. ((2015))***. Our study directly investigated the impact of rearing at 29 °C on different motor functions over the lifespan, and the cellular physiological links between proposed molecular mechanisms (***Harman, 1956; Finkel and Holbrook, 2000; Ziegler et al., 2015)*** and behavioral alterations previously described for *Sod* mutant flies.

We also present an overall assessment of the trajectory of neural circuit performance in the generation of different categories of motor behaviors along normal and altered lifespans. These include flight pattern generation (***Levine, 1973; Harcombe and Wyman, 1977)*** the giant fiber (GF) pathway-mediated jump and flight escape reflex (***Tanouye and Wyman, 1980; Gorczyca and Hall, 1984; Engel and Wu, 1992)***, habituation of the GF pathway escape response to repetitive stimulation (a form of non-associative learning, (***Engel and Wu, 1996, 1998))*** and a stereotypic seizure repertoire triggered by electroconvulsive stimulation (ECS, ***Pavlidis and Tanouye (1995)***; ***Kuebler and Tanouye (2000)***; ***Lee and Wu (2002***, 2006); ***Lee et al. ((2019))***. Our analysis has identified age-resilient and age-sensitive circuit properties that are attributable to changes in distinct circuit components. Furthermore, we found WT flies reared at 29 °C manifested accelerated trajectories of age-dependent alterations resembling those at 25 °C at a normalized timescale, while *Sod* mutants displayed distinctively altered pattern of alterations. Our findings also identify several neurophysiological parameters that could serve as quantitative functional indices for staging age-related decline. These findings have been reported in part in PhD dissertations (***Ruan, 2008; Iyengar, 2016)***.

## Results

### Flight Performance and DLM Firing Activity across the Lifespan: Effects of High-Temperature Rearing and *Sod* Mutation

The remarkable flight ability of *Drosophila* has been well documented (***Heisenberg and Wolf, 1984; Dickinson, 2014)***, with flights often lasting for hours in tethered flies (***Götz, 1968, 1987)***. To assess age-related changes in flight ability over the lifespan, we examined sustained tethered flight (Figure 1A, cf. ***Iyengar and Wu (2014))*** to delineate characterisic alterations in flight muscle electrical activity patterns and corresponding biomechanical parameters, and to reveal the effects of high-temperature and oxidative stressors on the aging process.

As established in *Drosophila* and other organisms, the aging process manifests a non-linear progression, indicated by varying rates of mortality at different stages in the lifespan (Figure 1B, cf. ***Vaupel et al. ((1998))***. We examined WT flies at 25 and 29 °C; two commonly used rearing temperatures in *Drosophila* aging studies. As previously reported, at 29 °C the fly lifespan was nearly halved as compared to 25 °C, a convenient feature frequently used to expedite aging studies (Figure 1B; ***Loeb and Northrop (1917)***; ***Leffelaar and Grigliatti (1983)***; ***Ruan (2008))***. At both temperatures, the lifespan trajectories follow an inverse sigmoidal curve, with “plateau”, “shoulder” and “tail” phases (***Curtsinger et al., 1992)***. We have also determined the effects of increased oxidative stress by characterizing *Sod* mutant flies, reared at a slightly lowered rearing temperature, 23 °C rather than 25 °C in order to expand the rather short *Sod* lifespan for more precise chronological tracking of data. Notably, the early plateau phase is absent in *Sod* lifespan curve at 23 °C (Figure 1B), retaining a distinctive feature in previously reported *Sod* life span curves obtained at higher temperatures (25 or 29 °C, ***Phillips et al. ((1989)***; ***Ruan (2008))***, which highlights the characteristic high mortality of young *Sod* flies.

**Figure 1.**
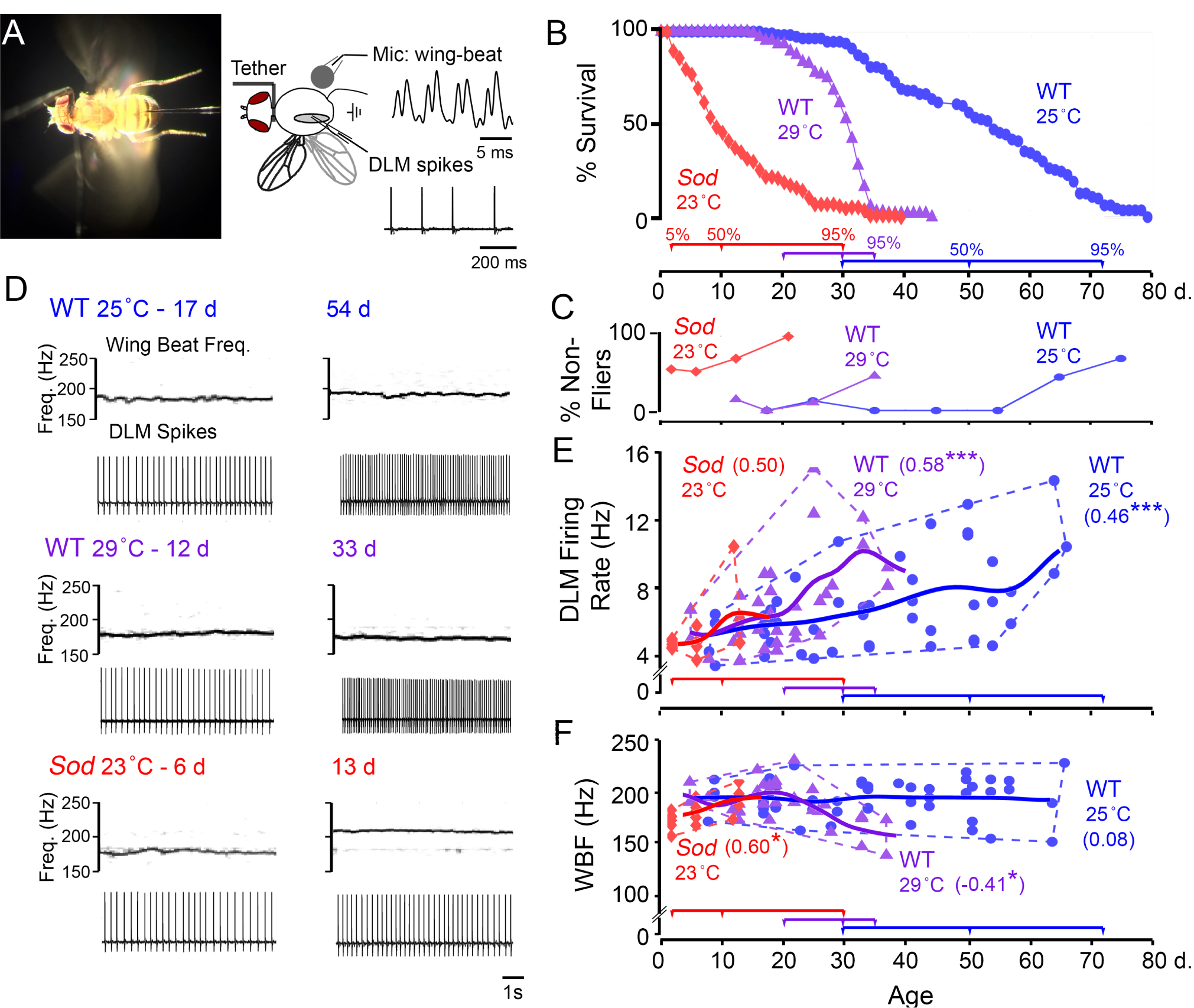
Flight motor performance across the lifespan of WT control and Sod mutant lies. (A, Left) Photograph of tethered fly preparation used in the flight and electrophysiological assays. Spiking activity in the DLM was monitored by a tungsten microelectrode, and acoustic signals generated by wing beats were picked up via a microphone placed below the fly. (Right) A schematic drawing of the preparation and sample waveforms of the respective signals. (B) Lifespan curves of WT flies (reared at 25 °C, blue circles, n = 110; and 29 °C, purple triangles, n = 104), and *Sod* mutants (red diamonds, reared at 23 °C, n = 134). Colored timelines below indicate the ages corresponding to 5%, 50% and 95% mortality for the respective curves. (C) The proportion of non-fliers (see Methods for definition) across the lifespan (n > 10 for each point). (D) Representative spectrograms of the microphone signal (upper panels) indicating the wing beat frequency (WBF) and corresponding traces of DLM spiking (lower panels) during sustained flight in younger (left column) and aged flies (right column) from the populations tested. (E-F) Scatterplots of the DLM firing rate (E) and WBF (F) during sustained flight in 25 °C- and 29 °C-reared WT flies (blue circles and purple triangles respectively), and in *Sod* mutants (red diamonds). Trend lines represent a Gaussian-kernel running average of the age-trajectory for the three populations (kernel window sizes of 10, 6 and 6 d respectively). The age-dependent Spearman’s rank correlation coefficient (r_s_) is indicated for each population (n = 48, 33, and 15 flies for WT 25 °C-, WT 29 °C-reared and *Sod* respectively). For this and following figures, * p < 0.05, ** p < 0.01, *** p < 0.001.

We found that flight ability was well-preserved in some aged WT flies reared at either 25 or 29 °C, displaying robust tethered flights beyond the 30-s sustained flight duration, a cut-off criterion which effectively distinguishes flight-defective mutants from WT individuals (***Iyengar and Wu, 2014)***. The “non-fliers” were mostly encountered in flies aged beyond the median lifespan (55 and 30 d for individuals reared at 25 and 29 °C respectively, Figure 1C). In striking contrast, many young *Sod* flies (∼50%) were not capable of sustained flight beyond 30 s, and this proportion further increased with age, with few *Sod* flies beyond the median lifespan (9 d) capable of flight beyond 30 s (Figure 1C).

The differences between WT and *Sod* mutants in flight ability prompted us to examine the wing beat frequency (WBF) as well as flight muscle DLM firing rate, across the lifespan. Using protocols established for the tethered flight preparation (***Iyengar and Wu, 2014***), we were able to acoustically monitor WBF via a microphone and simultaneously record DLM spiking activity (Figure 1A, D). The WBF is largely determined by the natural mechanical resonance frequency of the thorax case, which is powered by alternating activation of two sets of indirect flight muscles, DLMs and dorsalventral muscles or DVMs (***Chan and Dickinson, 1996)***. Isometric contraction of indirect flight muscles is stretch-activated and coincides with mechanical oscillation of the thorax, while their spiking activity, occurring at a much lower frequency than WBF, facilitates Ca^2+^ influx for force generation (***Dickinson and Tu, 1997; Lehmann and Dickinson, 1997***). Therefore, changes in muscle tension without shortening allow microelectrode recordings of the DLM spikes with minimal cell damage during prolonged flight.

Across the lifespan of WT flies, we noted a clear trend of increasing DLM firing rates which nearly doubled in the oldest flies examined (Figure 1D-E, Spearman rank correlation coefficient: r_s_ = 0.46, p = 9.9 × 10^−4^, and r_s_ = 0.58, p = 3.6 × 10^−4^, respectively, for 25 and 29 °C-reared flies). Accompanying this increasing trend, we noted an apparent overall increase in variability (Figure 1E, dashed distribution outline). In contrast to this age-associated variability in DLM firing, the WBF showed little change throughout lifespan for the fliers in the three populations. In 25 °C-reared WT flies both young and old individuals displayed average WBF of ∼190 Hz, with a narrower spread within a +/- 20 Hz (Figure 1F). We observed only a small, but detectable, reduction in frequency across the lifespan of 29 °C reared flies (r_s_ = −0.41, p = 0.19). Remarkably, the small population of *Sod* fliers displayed largely normal DLM firing rate and WBF with an indicative upward trend in both DLM firing rate and WBF (r_s_ = 0.50, p = 0.06 and r_s_ = 0.60, p = 0.018, respectively, Figure 1E-F). Taken together, our analysis of flight patterns suggests that the biomechanical properties of flight remained largely stable across the lifespan, while increases in DLM spiking mean frequency and variability reflected potential age-related alterations in central motor circuits.

In the following sections, we adopt the measure of “biological age”, by tracking the percent mortality of the population, in order to facilitate comparisons of the aging trajectories of functional alterations across different genotypes and conditions. This can be achieved by a re-normalization of the time axis to reflect the cumulative mortality based on the chronology lifespan curves (Figure 1B). As shown in the following figures, we collected data at different biological ages, for WT flies reared at 25 °C, biological ages at <1%, 5%, 50% and 95% mortality corresponded to the chronological ages of 7, 31, 50 and 72 days, whereas for 29 °C rearing, this shifted to 7, 20, 30 and 35 days. Since *Sod* flies lacked a prolonged initial plateau phase but showed a lingering “tail” in the lifespan curve, the biological ages targeted for data collection were modified to compress the first time interval to <5% mortality, and 70% mortality was added before the final 95% stage so as to allow sampling within the prolonged “tail”. The stages for *Sod* flies at <5%, 50%, 70%, and 95% mortality therefore corresponded to 2, 9, 14, and 30 days. The distinct profile of *Sod* aging trajectory indicates markedly different effects exerted by oxidative and high temperature stressors on the mortality rate along the lifespan. It is worth noting that despite the early precipitous drop and a shortened population medium lifespan, a small fraction of Sod mutant flies nevertheless exhibited relatively prolonged survivorship, giving rise to the characteristic, disproportionally prolonged tail in its lifespan curve.

### Age-Resilient and -Vulnerable Properties of the Giant Fiber (GF) Circuit that Mediates a Jump-and-Flight Escape Relex

One of the best-studied motor circuits in adult *Drosophila* is the giant fiber (GF) pathway that triggers a jump-and-flight escape reflex critical for survival (Figure 2A, cf. ***Tanouye and Wyman (1980)***; ***Trimarchi and Schneiderman (1995b)***; ***von Reyn et al. ((2017))***. Visual and other sensory inputs to the GF interneuron dendrites in the brain initiate the action potential propagating along the descending axon to drive a set of motor neurons in the thoracic ganglion. The DLM motor neuron receives the GF command signal indirectly via a peripherally synapsing interneuron (PSI) to evoke a DLM spike. This axo-axonal GF-PSI contact has been characterized as a mixed electrical and cholinergic synapse (***King and Wyman, 1980; Gorczyca and Hall, 1984; Blagburn et al., 1999; Allen and Murphey, 2007***). It has been well established that electric stimuli across the brain evoke two types of GF pathway-mediated DLM responses with distinct latencies (Figure 2B). Low-intensity stimulation across the brain recruits afferent inputs to the GF neuron, thereby triggering a relatively longer latency response (LLR, 4 - 6 ms, ***Elkins and Ganetzky (1990)***; ***Engel and Wu (1992***, 1996)), whereas high-intensity stimuli directly activate GF axonal spikes bypassing the brain afferents and resulting in a shorter latency response (SLR, 0.8 - 1.6 ms, ***Tanouye and Wyman (1980)***; ***Engel and Wu (1992))***.

**Figure 2.**
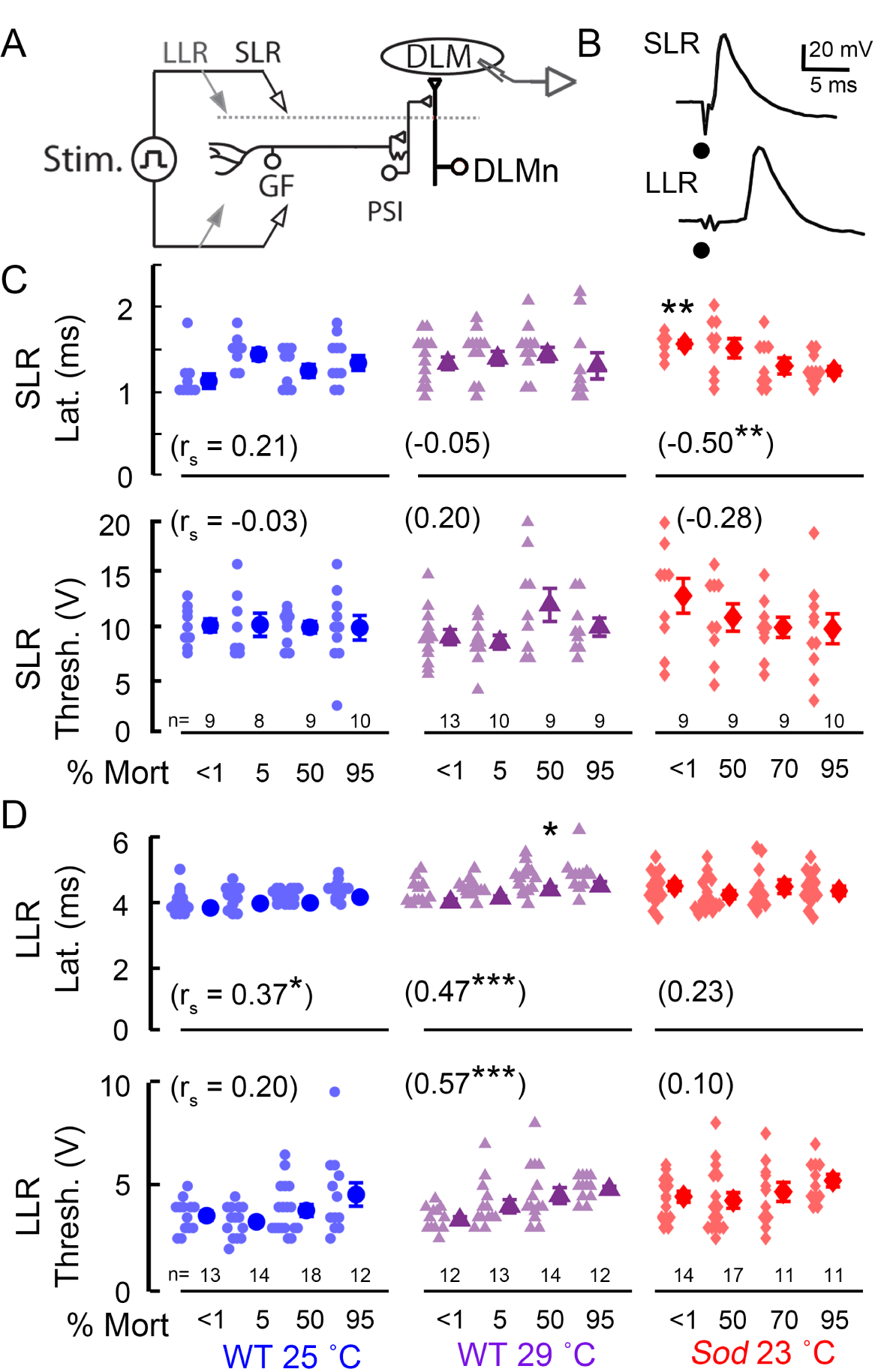
Performance characteristics of the GF pathway and its afferent inputs in aged WT and Sod mutant lies. (A) Overview of the GF “jump-and-flight” escape pathway. The GF neuron activates the indirect flight muscle DLM through a peripherally-synapsing interneuron (PSI) which innervates the DLM motor neuron (DLMn). For clarity, the GF branch that directly innervates the jump muscle (TTM) is not shown. (B) Electrical stimulation across the brain activates the GF pathway, and motor output is monitored via DLM action potentials in the tethered fly. High-intensity stimuli directly activate the GF neuron, giving rise to short-latency responses (SLRs). Lower-intensity stimuli recruit GF afferent inputs, resulting in the DLM long-latency responses (LLRs). Dots indicate stimulus artifacts. (C-D) SLR and LLR latency and threshold across the lifespan of WT and Sod flies. In this figure and in Figures 3 and 5, mean and SEM are indicated to the right of each dataset. Each data point represents measurement from a single fly. Asterisks indicate significant difference against mortality-matched, 25 °Creared WT flies (Kruskal-Wallis ANOVA, rank-sum post hoc test). The numbers in parenthesis indicate Spearman correlation coefficient (r_s_) of the parameter measurements vs. age categories.

We found clear distinctions between SLR and LLR in age-dependent modifications across the lifespan. Figure 2C shows robust maintenance of the basic properties (axonal conduction), of the GF neuron and downstream elements (PSI and DLM motor neurons) in aging WT flies, as indicated by the well preserved SLR parameters (see Table 1 for sample means, standard deviations and coefficients of variation). No significant age-dependent alterations were observed in the latency or threshold of the SLR for either 25 or 29 °C-reared WT flies, consistent with a previous report on well-maintained SLR properties up to a late stage of the lifespan (cf. ***Martinez et al. ((2007))***. Nevertheless, we did observe a significant change in SLR properties in young *Sod* mutants which displayed a retarded latency compared to 25 °C-reared WT counterparts (SLR latency mean: 1.6 vs 1.1 ms, p = 0.002, Kruskal-Wallis ANOVA, Figure 2C, Table 1). Furthermore, we noted an age-dependent decrease in this parameter for *Sod* (r_s_ = −0.50, p = 0.0015). This apparently paradoxical improvement brought the SLR latency parameters to a range directly comparable to those of WT flies, likely reflecting a progressive change in the Sod population composition, as healthier individuals persist and become better represented, while weaker ones die off.

**Table 1.**
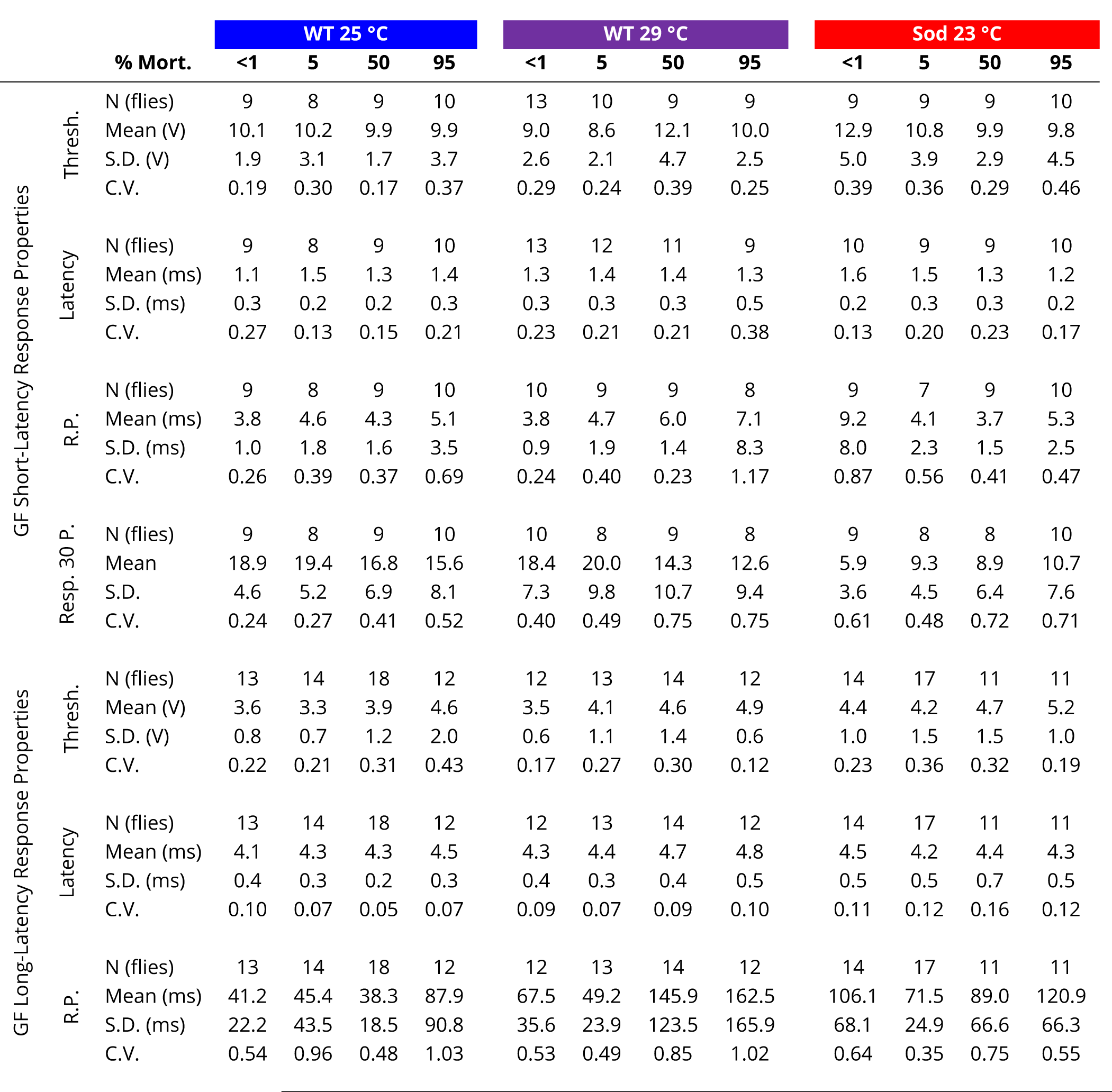
Features of GF Pathway SLR and LLR performance in young and aged flies

|  |  | % Mort. | WT 25 °C |  |  |  | WT 29 °C |  |  |  | Sod 23 °C |  |  |  |
| --- | --- | --- | --- | --- | --- | --- | --- | --- | --- | --- | --- | --- | --- | --- |
|  |  |  | <1 | 5 | 50 | 95 | <1 | 5 | 50 | 95 | <1 | 5 | 50 | 95 |
| GF Short-Latency Response Properties | Thresh. | N (flies) | 9 | 8 | 9 | 10 | 13 | 10 | 9 | 9 | 9 | 9 | 9 | 10 |
|  |  | Mean (V) | 10.1 | 10.2 | 9.9 | 9.9 | 9.0 | 8.6 | 12.1 | 10.0 | 12.9 | 10.8 | 9.9 | 9.8 |
|  |  | S.D. (V) | 1.9 | 3.1 | 1.7 | 3.7 | 2.6 | 2.1 | 4.7 | 2.5 | 5.0 | 3.9 | 2.9 | 4.5 |
|  |  | C.V. | 0.19 | 0.30 | 0.17 | 0.37 | 0.29 | 0.24 | 0.39 | 0.25 | 0.39 | 0.36 | 0.29 | 0.46 |
|  | Latency | N (flies) | 9 | 8 | 9 | 10 | 13 | 12 | 11 | 9 | 10 | 9 | 9 | 10 |
|  |  | Mean (ms) | 1.1 | 1.5 | 1.3 | 1.4 | 1.3 | 1.4 | 1.4 | 1.3 | 1.6 | 1.5 | 1.3 | 1.2 |
|  |  | S.D. (ms) | 0.3 | 0.2 | 0.2 | 0.3 | 0.3 | 0.3 | 0.3 | 0.5 | 0.2 | 0.3 | 0.3 | 0.2 |
|  |  | C.V. | 0.27 | 0.13 | 0.15 | 0.21 | 0.23 | 0.21 | 0.21 | 0.38 | 0.13 | 0.20 | 0.23 | 0.17 |
|  | R.P. | N (flies) | 9 | 8 | 9 | 10 | 10 | 9 | 9 | 8 | 9 | 7 | 9 | 10 |
|  |  | Mean (ms) | 3.8 | 4.6 | 4.3 | 5.1 | 3.8 | 4.7 | 6.0 | 7.1 | 9.2 | 4.1 | 3.7 | 5.3 |
|  |  | S.D. (ms) | 1.0 | 1.8 | 1.6 | 3.5 | 0.9 | 1.9 | 1.4 | 8.3 | 8.0 | 2.3 | 1.5 | 2.5 |
|  |  | C.V. | 0.26 | 0.39 | 0.37 | 0.69 | 0.24 | 0.40 | 0.23 | 1.17 | 0.87 | 0.56 | 0.41 | 0.47 |
|  | Resp. 30 P. | N (flies) | 9 | 8 | 9 | 10 | 10 | 8 | 9 | 8 | 9 | 8 | 8 | 10 |
|  |  | Mean | 18.9 | 19.4 | 16.8 | 15.6 | 18.4 | 20.0 | 14.3 | 12.6 | 5.9 | 9.3 | 8.9 | 10.7 |
|  |  | S.D. | 4.6 | 5.2 | 6.9 | 8.1 | 7.3 | 9.8 | 10.7 | 9.4 | 3.6 | 4.5 | 6.4 | 7.6 |
|  |  | C.V. | 0.24 | 0.27 | 0.41 | 0.52 | 0.40 | 0.49 | 0.75 | 0.75 | 0.61 | 0.48 | 0.72 | 0.71 |
| GF Long-Latency Response Properties | Thresh. | N (flies) | 13 | 14 | 18 | 12 | 12 | 13 | 14 | 12 | 14 | 17 | 11 | 11 |
|  |  | Mean (V) | 3.6 | 3.3 | 3.9 | 4.6 | 3.5 | 4.1 | 4.6 | 4.9 | 4.4 | 4.2 | 4.7 | 5.2 |
|  |  | S.D. (V) | 0.8 | 0.7 | 1.2 | 2.0 | 0.6 | 1.1 | 1.4 | 0.6 | 1.0 | 1.5 | 1.5 | 1.0 |
|  |  | C.V. | 0.22 | 0.21 | 0.31 | 0.43 | 0.17 | 0.27 | 0.30 | 0.12 | 0.23 | 0.36 | 0.32 | 0.19 |
|  | Latency | N (flies) | 13 | 14 | 18 | 12 | 12 | 13 | 14 | 12 | 14 | 17 | 11 | 11 |
|  |  | Mean (ms) | 4.1 | 4.3 | 4.3 | 4.5 | 4.3 | 4.4 | 4.7 | 4.8 | 4.5 | 4.2 | 4.4 | 4.3 |
|  |  | S.D. (ms) | 0.4 | 0.3 | 0.2 | 0.3 | 0.4 | 0.3 | 0.4 | 0.5 | 0.5 | 0.5 | 0.7 | 0.5 |
|  |  | C.V. | 0.10 | 0.07 | 0.05 | 0.07 | 0.09 | 0.07 | 0.09 | 0.10 | 0.11 | 0.12 | 0.16 | 0.12 |
|  | R.P. | N (flies) | 13 | 14 | 18 | 12 | 12 | 13 | 14 | 12 | 14 | 17 | 11 | 11 |
|  |  | Mean (ms) | 41.2 | 45.4 | 38.3 | 87.9 | 67.5 | 49.2 | 145.9 | 162.5 | 106.1 | 71.5 | 89.0 | 120.9 |
|  |  | S.D. (ms) | 22.2 | 43.5 | 18.5 | 90.8 | 35.6 | 23.9 | 123.5 | 165.9 | 68.1 | 24.9 | 66.6 | 66.3 |
|  |  | C.V. | 0.54 | 0.96 | 0.48 | 1.03 | 0.53 | 0.49 | 0.85 | 1.02 | 0.64 | 0.35 | 0.75 | 0.55 |

In contrast to the robust SLR properties, LLR parameters revealed clear changes in afferent inputs from higher centers to the GF neuron. Strikingly, among the oldest WT flies (e.g. 25 °C, 95% mortality), LLRs were not elicited in more than half of them (14 of 26), indicating severely compromised afferent inputs to the GF. Additionally, both 25 and 29 °C-reared WT populations displayed a small (< 10%), but detectable, age-dependent increases in LLR latency (r_s_ = 0.37 and 0.47; p = 0.003 and 0.0005, respectively, see also Table 1), and a modest age-dependent increase in LLR threshold (r_s_ = 0.57, p = 1.3 × 10^−5^) was found in 29 °C-reared flies. These relatively stable LLR properties were not grossly disrupted in the *Sod* mutant flies that remained responsive (Figure 2D).

In summary, our SLR results indicate that the GF and its downstream elements, including the PSI and DLM motor neurons, remain robust and could reliably transmit vital signals to evoke an escape reflex even in aged flies. However, the input elements upstream to the GF neuron showed more clear age-dependent functional decline, which could lead to a catastrophic collapse of LLR in some of the extremely old individuals (95% mortality) that were devoid of LLR.

### Alteration of SLR and LLR Refractory Period by High-Temperature and Oxidative Stressors

The robust, well-maintained basic properties of the GF and its downstream elements provide reliable readouts for examination of activity-dependent plasticity across the lifespan to uncover aging-vulnerable neural elements upstream of the GF neuron. By employing well-established stimulus protocols in the tethered fly preparation for activity-dependent processes, including paired-pulse refractory period, and habituation to repetitive stimulation (***Gorczyca and Hall, 1984; Engel and Wu, 1996***), we were able to establish modification due to aging in these use-dependent properties, as well as their alterations by high-temperature or oxidative stressors.

The refractory period (RP), for both SLR and LLRs, is defined as the shortest interval between paired stimulus pulses at which a second DLM response could be evoked (Figure 3A inset). We found WT flies across the lifespan displayed similar SLR RP performance when reared at 25 °C, while 29 °C-reared WT flies displayed an apparent, but not statistically significant trend of increasing RP over the lifespan. In a complimentary set of experiments, we delivered 30 SLR-evoking stimuli at 200 Hz and recorded the number successful responses (Resp. 30 p.). Across the lifespan, both 25 °C and 29 °C-reared flies displayed small alterations in performance, with changes not reaching statistical significance (Supplemental Figure 1). The refractoriness of LLR in older WT flies reared at 25 °C was similar to that of their younger counterparts. However, 29 °C-reared WT flies showed a significant age-dependent increase in LLR RP, most evident in the older populations (Figure 3B, 50 & 95% mortality). Notably, the distributions of SLR and LLR RPs in aged groups showed larger skews with several individuals displaying disproportionally lengthened refractory periods (less pronounced in SLR but most evident in LLR RP, Figure 3B). Furthermore, the oldest groups (95% mortality) consistently displayed the largest coefficients of variation of the LLR refractory periods datasets (Table 1). These observations highlight the stochastic nature of age-related functional deterioration in the upstream circuit elements afferent to the GF pathway.

**Figure 3.**
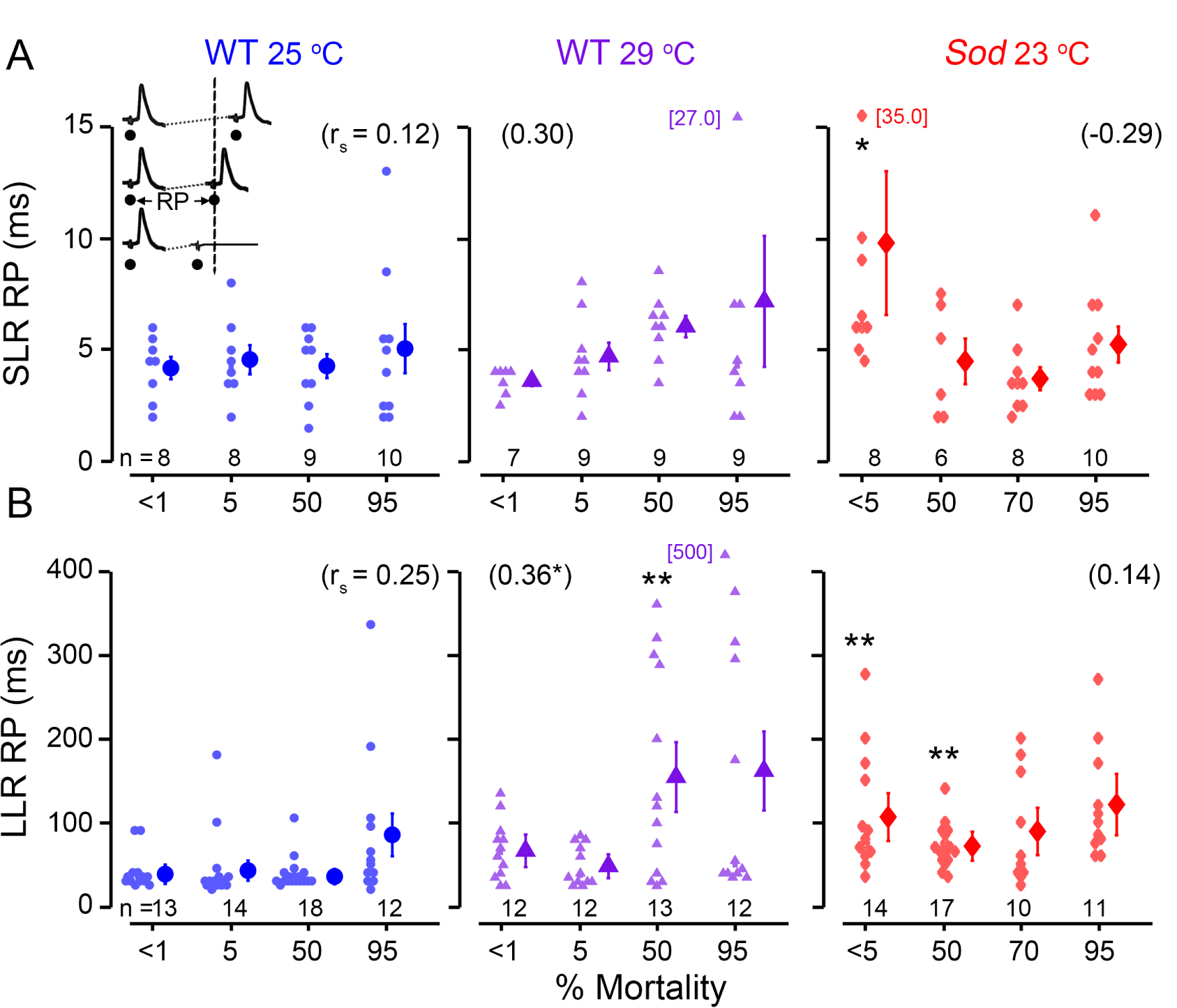
Paired-pulse refractory characteristics of GF-mediated DLM shortand long-latency responses. The paired-pulse refractory period (RP), represents the shortest inter-stimulus interval where the second stimulus evokes a DLM spike; shorter stimulus intervals fail to recruit two spikes (inset). (A-B) Scatter plots of the SLR (A) and LLR (B) RPs from WT flies reared at 25 °C and 29 °C as well as *Sod* flies reared at 23 °C are shown over the course of their lifespans, presented as percent mortality. Outlier values exceeding the axis range are indicated adjacently. See Figure 2 legend for statistical treatment and presentation.

Remarkably, young *Sod* flies represented the most heterogeneous population among different age groups in terms of refractory periods. Presumably the population included a portion of severely defective, very short-lived individuals that accounted for the initial drop and the diminished plateau phase of the lifespan curve (Figure 1B). Our observations revealed significant lengthening in both SLR and LLR refractory periods in very young (< 5% mortality) *Sod* mutants (SLRs p < 0.05, and LLRs, p < 0.01, Figure 3A-B, Kruskal-Wallis ANOVA, Bonferroni-corrected rank-sum post hoc test). Similarly, *Sod* mutants displayed significantly lower Resp. 30 p., indicating worse performance compared to WT counterparts (Supplemental Figure 1). In conjunction with a clear increase in SLR latency (Figure 2C), the RP lengthening point towards severe GF defects in a sub-population of young *Sod* flies. On the other hand, the surviving aged *Sod* mutants showed relatively stable SLR & LLR responses, with less substantial age-dependence beyond 50% mortality (Figure 3A-B).

### Aging Progression of Higher-Level Activity-Dependent Signal Processing: Acceleration of LLR Habituation over Lifespan

The above results suggest the functional integrity of the GF and its downstream elements as indicated by SLR performance (Figures 2 & 3) is age-resilient across the lifespan. Nevertheless, high-temperature stress and particularly *Sod* mutation can exert clear effects on activity-dependent aspects of GF-mediated DLM responses, as indicated by refractory period of LLRs (Figure 3B). In *Drosophila*, the jump-and-flight startle reflex shows habituation upon repetitive visual stimulation (***Trimarchi and Schneiderman, 1995a***), which is recapitulated in the habituation of LLRs (***Engel and Wu, 1996***). Habituation represents a non-associative form of learning that involves experience-dependent plasticity in higher functions (***Engel and Wu, 2009; Rankin et al., 2009)***.

To further investigate age-related changes in the plasticity of upstream processing of GF inputs, we examined habituation of LLRs (***Engel and Wu, 1996, 1998)***. Upon evoking LLRs repetitively, the DLM responses habituate, as subsequent stimuli fail to recruit LLRs (Figure 4A). However, LLRs may be rapidly restored following the application of a dis-habituation stimulus of a different modality, such as an air-puff, excluding sensory adaptation or motor fatigue as the basis for LLR failure (Figure 4A). We applied 100 LLR-evoking stimuli at three frequencies (1, 2 and 5 Hz), and adopted a habituation criterion as the number of stimuli required to reach 5 consecutive LLR failures (F5F, Figure 4A-B, cf. ***Engel and Wu (1996))***. The large number of stimuli used in this protocol provides greater power to resolve age-dependent alterations in an intrinsically stochastic process.

**Figure 4.**
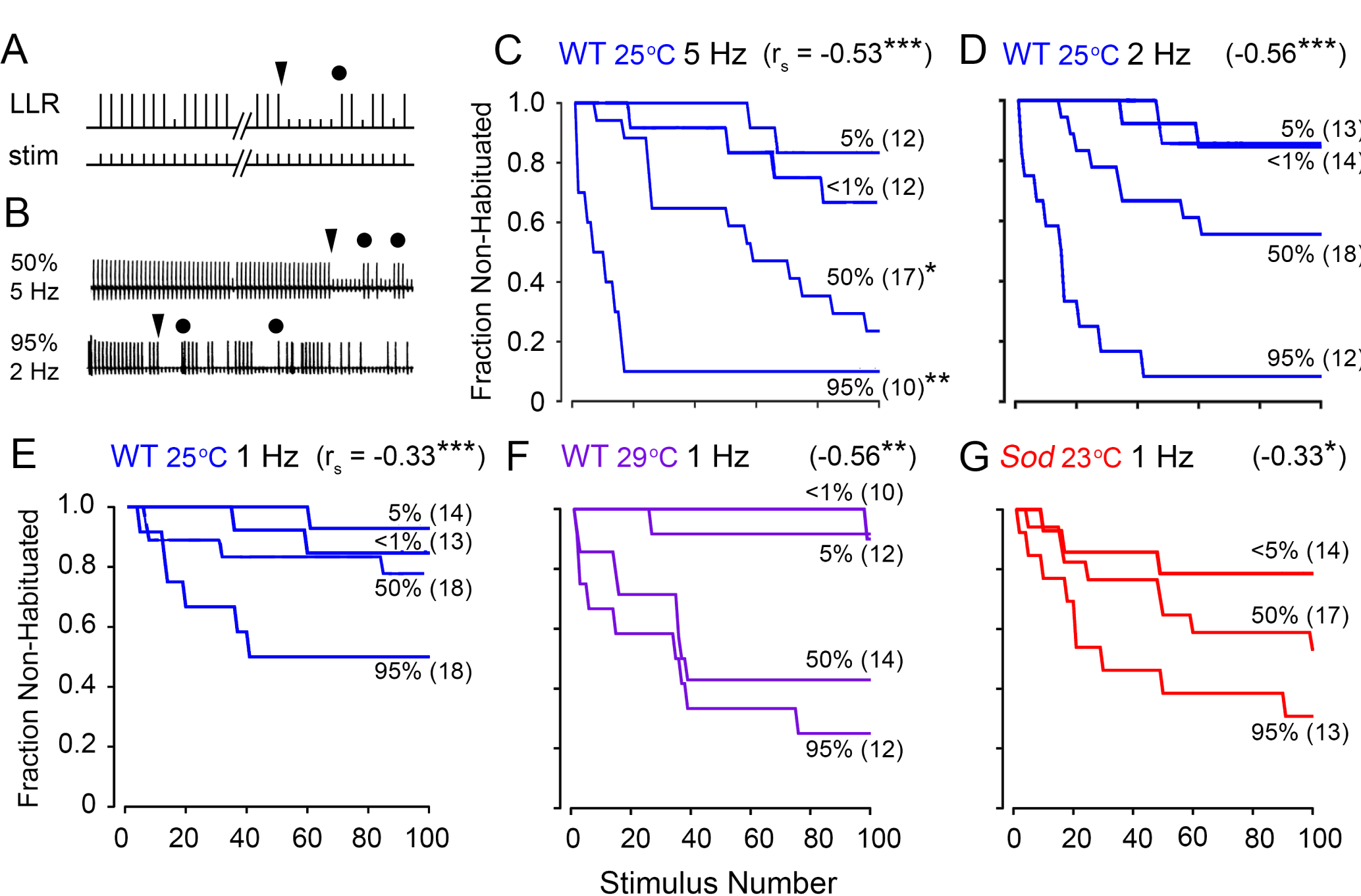
Age-dependent alterations in habituation kinetics of GF afferents. (A) Habituation of GF afferents. Habituation is operationally defined as the first instance of reaching five-consecutive DLM spike failures to LLR-evoking stimulation train of defined frequency (triangle,, cf. ***Engel and Wu (1996))***. Subsequent dis-habituation stimulus (circle, air puff) triggered recovery of LLRs, excluding sensory adaptation or motor fatigue as the basis for DLM response failure. (B) Sample traces from WT flies reared at 25 °C of slower habituation at 50 % mortality (upper trace, 5 Hz stimulation) and faster habituation at 95% mortality (lower trace, 2 Hz stimulation). (C-E) Frequency dependence of LLR habituation in 25 °C-reared WT flies to stimulation at 5 Hz (C), 2 Hz (D), and 1 Hz (E). The fraction of flies in the sample population of each age group (<1, 5, 50, 95% mortality) that were not habituated after a given number of stimuli is plotted. (F-G) Habituation in WT 29 °C-reared flies (F) and in *Sod* mutants (G) to 1 Hz stimulation. Sample sizes are indicated next to each plot. Asterisks at the end of the trace indicate significantly faster habituation compared to habituation in mortality-matched WT 25 °C-reared flies to 1 Hz stimulation (log-rank test). The age-dependent Spearman correlation coefficient (r_s_) is indicated in the upper-right corners of each panel.

Across the three stimulation frequencies examined (1, 2 and 5 Hz) for WT flies reared at 25 °C, we found that between < 1% and 5% mortality, during the plateau portion of the lifespan, the LLR habituation rate was largely unaltered with a large fraction of flies showing little habituation throughout the stimulation train (Figure 4C-E). However, during the mortality phase of the lifespan curve, 50% mortality and beyond, we observed a marked acceleration in LLR habituation, with the oldest flies habituating the fastest (r_s_ between −0.33 to −0.56, p < 0.001). As expected, we found more evident acceleration in habituation rate at higher stimulation frequencies (Figure 4C-E), which was most pronounced in the older populations (50 & 95 % mortality) examined. Together, our findings suggest that the brain circuits afferent to the GF pathway remained largely unaltered during the plateau portion of the lifespan curve, a substantial fraction of the lifespan. Nevertheless, across the lifespan, we could delineate the distinct habituation rates associated with the 5%, 50% and 95% age groups, particularly at the higher simulation frequency of 5 Hz, offering a quantifiable index of aging progression.

High-temperature rearing and *Sod* mutations further accelerated habituation. Compared to 25 °C-reared counterparts, 29 °C-reared WT flies displayed considerably faster habituation to 1 Hz stimulation in aged groups (50 and 95 % mortality, Figure 4E vs. F). Furthermore, *Sod* mutants (at 23 °C) showed an accelerated habituation rates compared to WT counterparts even in younger flies (< 5% mortality, Figure 4G vs, E & F), presumably correlated with the pronounced increase in LLR paired pulse refractory period in this age group (Figure 3B). Therefore, among the temperature- or oxidation-stressed groups, the habituation rate again can serve as a physiological marker of age progression even at low frequencies (∼1 Hz) of repetitive stimulation.

### Increasing Susceptibility to Electroconvulsive Induction of Seizures across the Lifespan

In addition to acting as an output of the flight pattern generator and GF pathway-mediated escape reflex described above, DLM action potentials may serve as reliable indicators of CNS spike discharges associated with seizures (***Pavlidis and Tanouye, 1995; Lee and Wu, 2002***). Such seizure episodes may reflect aberrant spiking originating from motor pattern generators located in the thorax, such as flight (***Harcombe and Wyman, 1977; Iyengar and Wu, 2014)*** and grooming (***Engel and Wu, 1992; Lee et al., 2019***). As shown in Figure 5A, high-frequency stimulation across the brain (200-Hz, 2-s train of 0.1-ms pulses, ***Lee and Wu (2002))*** can trigger a highly stereotypic electroconvulsive seizure (ECS) discharge repertoire, consisting of an initial discharge (ID), a period of paralysis reflected by GF pathway failure (F, in response to 1-Hz test pulses), and a second delayed discharge (DD) of spikes. This experimental protocol provides several stable and quantitative measures, within individual flies, suitable for assessing age-related modifications of CNS function, including the threshold to induce seizures (by adjusting intensity from 15 to 80 V, Figure 5B) and the discharge duration of DD spike trains (Figure 5C).

**Figure 5.**
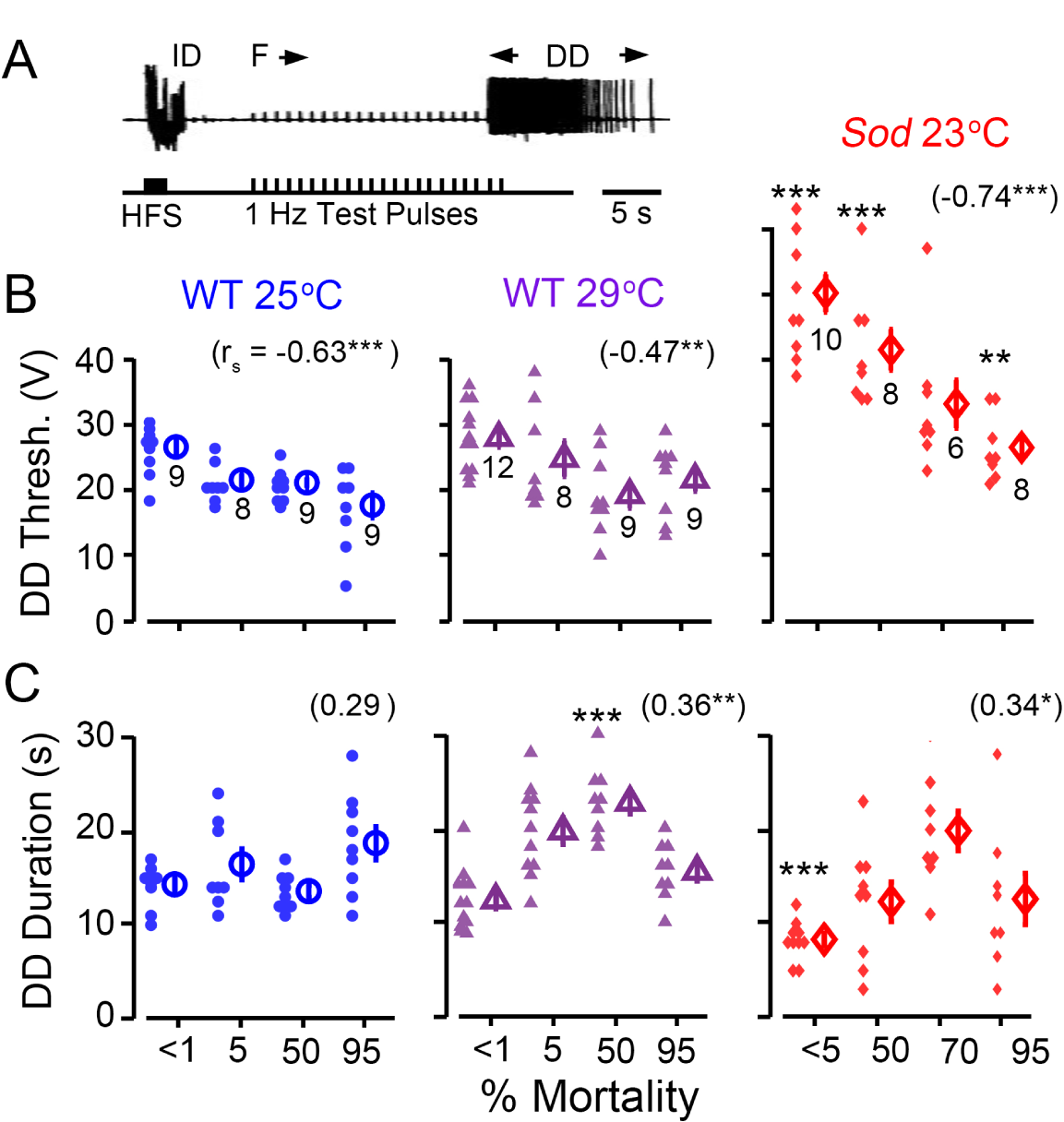
Decreasing threshold of electroconvulsive seizure (ECS) discharges in aging lies. (A) ECS activity was evoked by a short train (2 s) of high intensity, high frequency stimulation (HFS, 0.1 ms pulse duration, 200 Hz) across the head in tethered flies. DLM spike discharges serve as a convenient monitor of these seizures, and the pattern of these discharges is highly stereotypic (cf. ***Lee and Wu (2002))***, consisting of an initial spike discharge (ID), followed by a period of paralysis corresponding to failure of test pulse-evoked GF pathway responses (F), and a delayed spike discharge (DD). The example trace here shows the sequence in a young 25 °C-reared WT fly. (B-C) Scatterplots of the stimulation threshold to induce DD (B), and duration of the evoked DD (C) across the lifespans of 25 °C- and 29 °C-reared WT and *Sod* mutants. Mean and SEM are indicated to the right of each group, sample sizes below. See Figure 2 legend for other details of statistical treatment and presentation.

We found a clear trend of increasing seizure susceptibility with age in WT and *Sod* flies (Figure 5B). WT flies reared either at 25 or 29 °C displayed a substantial reduction in the DD threshold across their lifespan (−33% and −23% respectively for <1% vs. 95% mortality). In *Sod* mutants, the trend was even more pronounced (−46 % for 5 % vs. 95 % mortality). The effects of the *Sod* mutation were most striking in the youngest age group. In these flies (< 5% mortality), threshold was significantly higher than WT counterparts (p < 0.001, Kruskal-Wallis ANOVA, rank-sum post hoc), corroborating the other observations of extreme physiological defects found in this age group (Figures 2 & 3). Furthermore, this increase in ECS threshold was present across the *Sod* lifespan (p < 0.05, Kruskal-Wallis ANOVA, rank-sum post hoc) suggesting that elevated oxidative stress could render seizure-prone circuits less excitable.

Despite individual variability within each fly group, significant correlation coefficients suggest an enhanced excitability of seizure-prone circuits in aged flies (Figure 5B). Compared with other potential indicators, e.g. habituation rate (Figure 4), this age-dependent monotonic decrease of ECS threshold could provide a tighter, more quantitative measure of aging progression.

In contrast to the decreasing trend of seizure threshold over lifespan, there was a milder trend of increasing DD duration in WT flies reared at 29 °C, or in *Sod* mutants, but ending with a substantial drop in the oldest (95%) populations (Figure 5C, r_s_ = 0.36, p = 0.0041; and 0.34, p = 0.047 for WT 29 °C and *Sod* respectively). However, there was no clear age-dependent trend of increasing DD duration in 25 °C-reared WT flies (Figure 5C). In a complementary data set acquired at higher temporal resolution to resolve individual spikes, we examined the total number of spikes during the DD in these populations. Mirroring the DD duration observations, we found that 25 °C-reared WT flies did not display an age-dependent trend, while WT flies reared at 29 °C, as well as *Sod* mutants, showed modest increases in the DD spike count (Supplemental Figure 2, r_s_ = 0.50, p = 1.4 × 10^5^ for WT 29 °C; r_s_ = 0.27, p =0.04 for *Sod* mutants).

### Age-Trajectories of ID Firing Patterns Reveal a Potential Predictor of Mortality Rate

Alterations in ECS threshold and DD duration across lifespan (Figure 5) call for detailed characterization of ID spiking to extract additional salient features associated with aging progression. Since ID occurs as a brief burst (1 – 4 s, see Figures 5A and 6A) of high-frequency spiking activity, a set of ECS data at a higher temporal resolution enabled an analysis of ID spiking dynamics at the level of individual spikes (Figure 6A). Immediately following electroconvulsive stimulation, the ID spike patterns manifested striking, but more complex, age dependence and vulnerability to oxidative stress.

**Figure 6.**
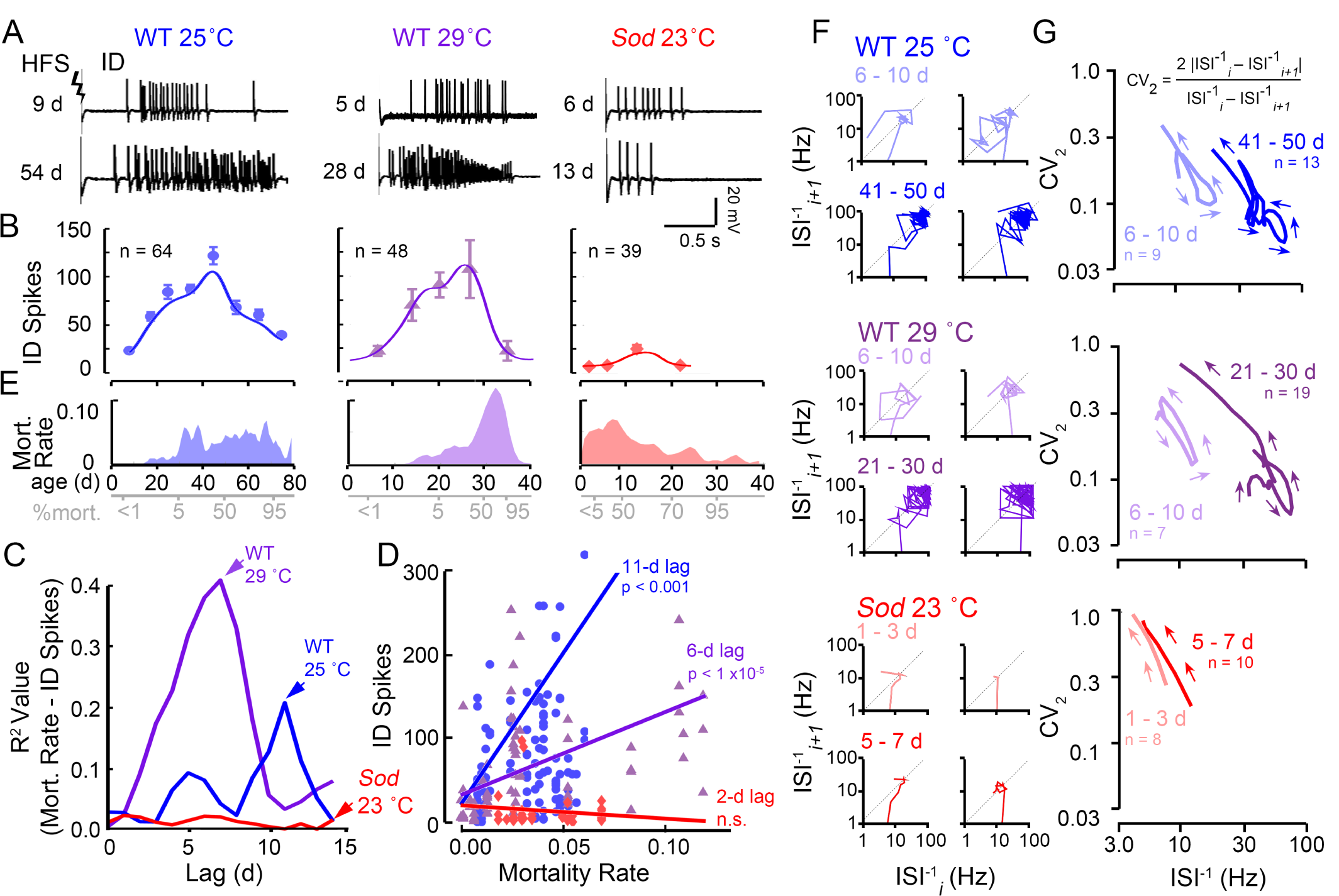
Age-dependent alterations in spike patterning of initial seizure discharge. (A) Representative high frequency stimulation-induced IDs from young (< 1% mortality; 9, 5 and 6 d traces) and aged (50% mortality; 54, 28 and 13 d traces) WT and *Sod* flies (stimulation voltage: 80 V). (B) Plot of the number of spikes within the ID across the lifespan (mean ± SEM for different age groups, and sample sizes across the populations are indicated). Trend lines represent a Gaussian-kernel running average (see Figure 1) of the age-trajectory for the three populations. (C) Plot of the mortality rates (defined as the negative slope of the lifespan curve, Figure 1B) across the lifespan for the three populations examined. (D) Linear cross-correlation between the mortality rate and ID spike count upon introducing a lag of 0 – 15 days to determine the optimal lag for maximal correlation, i.e. 11 and 6 d for 25 °C and 29 °C-reared WT flies. In *Sod* mutants no significant correlation between mortality rate and ID spike count was detected. (E) Plot of best correlation between of lagged ID spike count versus mortality rate for the three populations. (p-values as indicated). (F) Poincaré plots of representative IDs from young (< 1% mortality) and aged (50% mortality) flies. The instantaneous firing frequency of each inter-spike interval 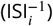 is plotted against the next spike interval 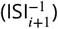 on a log scale (see text). (G) Averaged trajectories (as constructed in ***Lee et al. ((2019))*** of the inter-spike interval (ISI^−1^) versus the instantaneous coefficient of variation (CV_2_, as defined by Holt et al., 1996, see text). Lower values of CV_2_ indicate more rhythmic firing. Number of trajectories used to construct the average is as indicated. Note the similar changes in aged 25 °C and 29 °C–reared WT flies compared to young counterparts and distinctions between WT and *Sod* mutant discharges.

In younger WT flies (reared at either 25 or 29 °C), ID spike counts progressively increased with age until about 50% mortality (Figure 6A and B). Beyond 50% mortality, a decline in the ID spike count became apparent; notably the oldest flies (95% mortality) regressed to a spike count similar to those of young flies (< 1% mortality). In contrast to nearly 10-fold range for ID spike counts across the WT lifespan, *Sod* mutants displayed only rather sparse spiking, too few to shape a prominent age-related profile in their counts (Figure 6A and B). This observation mirrors the particular vulnerability of ECS discharges to oxidative stress, along with the drastically elevated threshold levels in ECS induction in *Sod* mutant flies (Figure 5B).

We noted that the curve of average ID spike count across the lifespan showed a gross resemblance in overall profile with the mortality rate curve. The progression of spike count changes apparently preceded the mortality rate changes in WT (compare Figure 6B and C). We sought an appropriate temporal shift between the ID spike count and mortality rate curves that would optimize regression fit. The iterations yielded the best correlation between the ID spike count and mortality rate with 11- and 6-d lags for WT flies reared at 25 and 29 °C, respectively (Figure 6D and E). These observations suggest that the ID spike production over the lifespan may serve a predictor for the ensuing mortality rate in WT flies. In other words, along with other physiological indicators for aging progression examined above (Figures 4 and 5B), ID spike counts might also provide a gross forecast of the mortality rate, which is the derivative of the lifespan curve. However, this relationship may not be applicable to a variety of genotypes since it does not seem to hold true for *Sod* flies (Figure 6A and B).

### Signatures of ID Spiking Patterns in Young and Old WT and *Sod* Flies

Salient features of spike patterns can be uncovered by systematical tracking of the successive changes in consecutive spike intervals to depict the spike train dynamics, as shown in the Poincaré plots (Figure 6F, cf. ***Lee et al. ((2019))***. In these plots, instantaneous firing frequency, defined as the inverse of an inter-spike interval (ISI^−1^), was first determined for each of the successive spike intervals in a spike train. Each ISI^−1^ was then plotted against the ISI^−1^ of the following spike interval and the process was reiterated sequentially for each spike in the ID spike train (i.e. ISI^−1^ vs. ISI^−1^ _*i*+1_ for the *i*^th^ spike in the sequence). For strictly periodic spiking, all data points of the Poincaré trajectory (PT) would fall at a single location along the diagonal. Irregular deviations from rhythmicity between successive spike intervals appear as perpendicular shifts away from the diagonal. Thus, the trajectory depicts the evolution of spiking frequency, while excursion patterns about the diagonal in the PT indicate characteristics of recurrent firing patterns. Poincaré plots of representative individual spike trains are shown in Figure 6F that readily capture the larger number of spikes with increasing ISI^−1^ in aged WT flies compared to younger counterparts (up to 100 Hz vs. < 30 Hz). In contrast, the few ID spikes in *Sod* flies, regardless of age, occurred in a lower frequency range (10 Hz).

To integrate information embedded in individual PTs to indicate overall differences in ID spiking dynamics between age groups, we replotted the PTs to carry both frequency and local variation for successive ISIs in order to combine them into an “average” trajectory (Figure 6G, see Methods and ***Lee et al. ((2019)*** for computational details). Briefly, for each ISI in the ID spike sequence, the ISI^−1^ was plotted against the instantaneous coefficient of variation (CV_2_), defined as: 2 | ISI^−1^ – ISI^−1^ _*i* +1_| / [ISI^−1^ + ISI^−1^ _*i* +1_] (***Holt et al., 1996***). Along the trajectory, low CV_2_ values correspond to more regular firing (lower fractional changes among adjacent spike intervals), while high CV_2_ values indicate where more abrupt changes in spike intervals occur. The IDs of young WT flies (< 1% mortality) reared at either 25 °C or 29 °C displayed rather similar ISI^−1^ vs CV_2_ trajectories compared to each other, with peak ISI^−1^ between 10 and 20 Hz and a similar range of CV_2_ in the spiking procession. Remarkably, for both rearing temperatures, the trajectories of aged WT flies displayed similar transformation from their young counterparts, with more complex, right-shifted trajectories with peak ISI^−1^s > 70 Hz (Figure 6G, 41 – 50 d and 21-30 d, respectively, for 25 °C and 29 °C-rearing), suggesting that the changes in circuit activity patterns across lifespan are conserved during high-temperature rearing. Nevertheless, the simple ISI^−1^ vs CV_2_ trajectories of both young and aged *Sod* stood in sharp contrast to WT trajectories, providing another major distinction between the effects of high-temperature and oxidative stressors and underscoring the hypoexcitable nature of the *Sod* mutant.

Clearly, these “average” trajectories present concisely the overall characteristics of the ID spiking activity, with precise temporal information from the initiation to cessation of the spike train. They are effective signatures for IDs of different age groups in each genotype; WT flies reared at 25 and 29 °C traverse similar terrains in the span of ID spike trains whereas *Sod* flies of various ages yield only abbreviated signatures, marking very limited terrains in specific regions in the frequency-variance landscape.

## Discussion

This report summarizes our initial efforts towards a quantitative description of neurophysiological changes during aging in several well-established motor circuits in *Drosophila*. Aging studies in flies have largely focused on how the lifespan is influenced by genetic modifications in cellular pathway signaling, such as those regulated by insulin-like peptide (e.g. ***Clancy et al. ((2001)***; ***Hwangbo et al. ((2004))*** target of rapamycin (TOR, e.g. ***Kapahi et al. ((2004))***, and oxidative stress (e.g. ***Tower (1996)***; ***Phillips et al. ((2000))***, and by environmental factors, including diet (e.g. ***Mair et al. ((2003)***; ***Piper and Partridge (2007)***; ***Skorupa et al. ((2008))*** mating experience (***Kuo et al., 2012; Gendron et al., 2014; Harvanek et al., 2017***), exercise (***Piazza et al., 2009; Sujkowski et al., 2015***), and trauma (***Katzenberger et al*., *2013*, ***2015)*****. In parallel, several established behavioral assays have provided readouts of age-related functional decline, including fast phototaxis and negative-geotaxis (e.g. ***Miquel et al. ((1976)***; ***Arking and Wells (1990)***; ***Simon et al. ((2006****), flight (****Miller et al., 2008***), memory (***Tamura et al*., *2003;*** ***Yamazaki et al., 2007***), courtship (***Cook and Cook, 1975; Partridge et al., 1987***), as well as sleep and other circadian activity patterns (***Koh et al., 2006; Bushey et al., 2010***). However, behavioral studies registering concomitant changes in neurophysiological processes remain to be explored in *Drosophila*, currently with only limited examinations focusing on electroretinogram (***Ueda et al., 2018***) or GF pathway SLR properties (***Martinez et al., 2007; Augustin et al., 2018***, ***2019; Blagburn, 2020***).

The readily quantifiable but distinct spiking activities of the DLM motor neuron driven by reflex commands and different motor pattern generators, including flight, GF-mediated escape and seizure discharges, have enabled us to characterize trajectories of age-related functional modifications in different *Drosophila* motor circuits. Quantitative comparisons revealed drastically different effects of high-temperature rearing and oxidative stress on the aging trajectories of the various neurophysiological parameters in WT and mutant flies.

### Aging-Resilient and -Vulnerable Aspects of Motor Circuit Function in *Drosophila*

In this study, we adopted a normalization in time scales of aging progression in accordance with % mortality (***Jones et al., 2014***) to facilitate comparisons across the genotypes and conditions that yield drastically different chronological lifespan curves (Figure 1B). A graphical overview emerges when comparing the age-trajectories of the various motor circuit parameters in different fly populations. Relative to the starting point (i.e. youngest age group), changes (in fold on a semi-logarithmic log_2_ scale) in various circuit functions can be compared directly for their relative vulnerability or resilience to aging (Figure 7).

**Figure 7.**
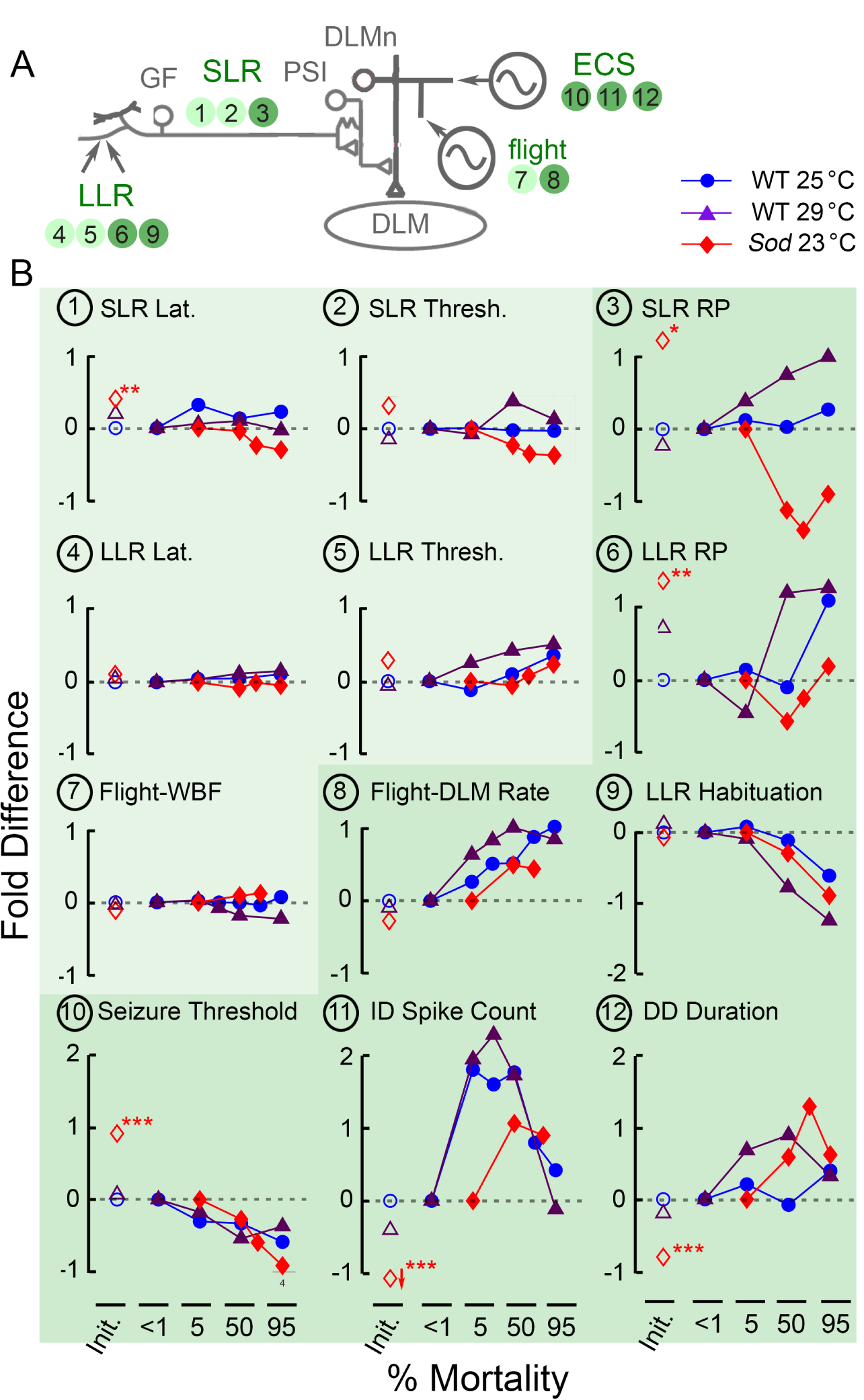
Distinct age-trajectories of different motor circuit properties examined in this study. (A) Schematic diagram of the circuit components generating motor patterns driving DLM activity examined in this study. Corresponding parameters displayed in panels in (B) are indicated by corresponding panel numbers. Darker backgrounds indicate that significant age-dependent alterations were detected. (B) Normalized changes in neurophysiological parameters of the respective motor circuits are plotted as function of % mortality. Changes are presented in terms of ‘fold change’ from the initial (<1% mortality) value (i.e. log_2_ [value/initial value]), such that +1 indicates doubling, while −1 indicates halving of the parameter’s value. First, the plot at the origin (open symbols, <1% mortality) indicate fold changes compared to young WT 25 °C for WT 29 °C and *Sod* 23 °C young flies. Second, the age-trajectories comparing flies of different ages to the youngest flies are indicate in fold changes (filled symbols: blue, purple and red for WT 25 °C, WT 29 °C, and *Sod* 23 °C respectively). Panels representing circuit parameters that remain relatively robust across the lifespan in all three populations are shaded in lightly shaded, while parameters that display clear age-dependent alterations in at least one rearing condition are shaded darker.

Our results suggest the operation and integrity of motor circuits fall into two general categories: aging-resilient (Figure 7, lighter shading) and -vulnerable (darker shading). Notably, the elements in the GF pathway that mediate the escape reflex appear more robust and show less functional decline up to the very end of the lifespan and display milder effects of high-temperature stress or ROS scavenger disruption, with little impact on SLR and LLR threshold and latency (Figure 7B-1, -2, -4 and -5). In contrast, substantial changes were seen in the more plastic, use-dependent, functions of these circuit components, including the twin-pulse refractory period of the GF pathway and the habituation of its afferent inputs (Figure 7B-3, -6, and -9). Furthermore, motor program-related DLM spike readouts, such as the flight and ECS discharge patterns, showed remarkable age-related modifications (Fig 7B-8, -10, -11 and -12).

### GF Pathway SLRs: GF Neuron and Downstream Elements

The GF pathway with its well-described neural elements and transmission properties offers an ideal model system to explore cellular mechanisms of aging and neurodegenerative diseases, as previously reported for SLR properties (***Martinez et al., 2007; Sofola et al., 2010; Zhao et al., 2010; Augustin et al., 2018***). Consistently, our characterization of SLRs underlying the escape reflex demonstrates well maintained basic mechanisms, for signal propagation from the GF neuron through the PSI and DLM motor neuron to postsynaptic DLMs. Over a large portion of the lifespan, including the oldest flies (95% mortality), WT flies reared at both room 25 and 29 °C did not show a clear trend of alteration in either SLR latency or threshold, but a decreasing trend in both threshold and latency was noted in *Sod* mutant flies, indicating a consequence of chronic oxidative stress (Figure 2C, Figure 7B-1 and -2).

More evident aging-vulnerable alterations were seen in the activity-dependent properties for flies grown under high-temperature or ROS stressors. For WT reared at 25 °C, frequency-dependent characteristics of the SLR, including the twin-pulse RP and Resp. 30 p, were relatively stable with no statistically significant age-dependent changes detected (Figure 3A and Table 1). However, this is not the case for 29 °C-reared WT or *Sod* mutant flies since detectable but opposite trends of changes were observed in these two categories of flies in both twin-pulse RP and Resp. 30 p. readouts (Figure 3A, Figure 7B-3, Table 1), suggesting differential vulnerable targets of these two types of stressors.

### GF Afferent Processing: LLR Refractory Period and Habituation

The processing of afferent signals from upstream circuits activates the GF pathway LLRs (Figure 7A). Similar to SLRs, we found activity-dependent properties of LLRs, i.e. RP and habituation were more markedly modified over the lifespan than the subtle changes in latencies and thresholds (Figures 2D, 3B, 4; Table 1; Figure 7B-4 and -5). Given the apparent stability of transmission of the GF neuron and downstream elements, these changes hint at age-dependent plasticity in GF afferents in the brain (***Engel and Wu, 1996***, ***1998)***, perhaps in identified visual (***Ache et al., 2019***) or mechanosensory ***Lehnert et al. ((2013)***; ***Mu et al. ((2014)***; ***Blagburn (2020)*** inputs.

We observed increasing trends of LLR latency and threshold as flies age (Figures 2D; Table 1; Figure 7B-4 and -5). However, even though age-dependent alterations observed in LLR latency are statistically significant, they are relatively minor in magnitude, constituting only small fractions in time scale within a 5-20 ms behavioral repertoire for visually or olfactorily triggered GF-mediated escape responses (***Engel and Wu, 1992; Trimarchi and Schneiderman, 1995a; Engel and Wu, 1996; von Reyn et al., 2017***).

We also noted that the LLR RP determined by the paired-pulse protocol revealed substantial heterogeneity in the older WT populations examined as indicated by extremely large spread of data points (50 and 95% mortality 25 and 29 °C, Figure 3B, Table 1), presumably a reflection of the stochastic nature of aging progression in the processing of GF afferent signals.

Importantly, the LLR habituation protocol (***Engel and Wu, 1996***, ***1998)*** demonstrated the clearest monotonic age-dependent changes in the performance of the GF pathway. In both 25 and 29 °C-reared WT, as well as *Sod* populations, aged flies habituated much faster compared to younger counterparts (Figure 4, Figure 7B-9). Importantly, across all populations, these effects were robust for all frequencies tested (1, 2, and 5 Hz, with the amplitude of r_s_ coefficients ranging from 0.33 to 0.56 with significant p values). Habituation is commonly considered a simple form of nonassociative learning (***Engel and Wu, 2009; Rankin et al., 2009***). Our study demonstrated that the *Drosophila* GF habituation rate increases monotonically and robustly with age (Figures 4 and 7B-9), rendering this plasticity parameter a potential quantitative index of aging progression for an important aspect of nervous system function.

### Flight

Throughout most of their lifespan, WT flies maintained their flight ability (Figure 1C), with stable WBFs (Figure 1D, F; Figure 7B-7) suggesting effectively adjusted biomechanical properties of flight muscles and thoracic case. Our findings are largely consistent with a previous report (***Miller et al., 2008***) where the WBF was stable beyond the median lifespan despite marked changes in muscle structure and stiffness (see also ***Chaturvedi et al. ((2019))***. Despite the nearly invariant WBFs, we found the motor circuit driving DLM firing during flight displayed a clear and largely monotonic increase in firing rate with age (Figure 1E, Figure 7B-8), highlighting a robust phenotypic change in both 25 °C and 29 °C-reared flies. Conceivably, the age-dependent increases in firing rate could indicate changes in motoneuron excitability or flight circuit signal processing among the relevant thoracic interneurons, which may reflect compensatory mechanisms to ensure flight ability, i.e increasing spiking to retain appropriate intracellular Ca^2+^ levels for powering sustained flight in aging myogenic stretch-activated DLMs (***Pringle, 1978; Gordon and Dickinson, 2006; Lehmann et al., 2013***).

### Electroconvulsive Seizure

In tethered flies, high frequency stimulation across the head recruits a remarkably stereotypic pattern of nervous system-wide spike discharges accompanied by wing-buzzes (in a fixed succession of ID, paralysis, and DD, cf. ***Lee and Wu (2002))*** reminiscent of the sequence of seizures and paralysis induced by mechanical shock in “bang-sensitive” mutants (***Ganetzky and Wu, 1982*; *Burg and Wu, 2012***). Importantly, the spiking associated with the ID and DD is independent of the GF system; during the ECS repertoire, transmission along the escape pathway is blocked (***Pavlidis and*** *Tanouye, 1995*). Analysis of decapitated flies and Ring-X gynandromorph mosaics has further indicated both ID and DD pattern generators reside in the thorax, although globally synchronized episodes of electric activities and quiescence concurrent with ID, paralysis and DD also emerge in different parts of the CNS (***Lee and Wu, 2002***). However, the ID and DD are likely derived from separable network origins, as they are independently vulnerable to different ion channel mutations (***Lee and Wu, 2006; Kasuya et al., 2019***).

In both WT and *Sod* flies, we noted a clear age-dependent decrease in ECS-evoked seizure threshold across the lifespan (Figure 5, Figure 7B-10). The monotonic characteristic of this reduction suggests seizure threshold may be considered as another quantitative index of the age progression in nervous system function. Indeed, in rats, aging reportedly reduced the threshold for kainate-induced seizures (***Liang et al., 2007***), and aged humans display increased incidence of seizure disorders (***Tallis et al., 1991; Leppik et al., 2006***).

ECS discharge spiking patterns appeared to be age-dependent as well (Figure 7B-11 and -12), most dramatically during the ID phase (Figure 6). In contrast to a monotonic trend, ID spiking increases initially with age, peaking around 50% mortality, then declines to earlier levels towards the end of the lifespan (Figures 6B and 7B-11). Our finding that changing ID spiking intensity seemingly provides a forecast for subsequent mortality rate is intriguing. Indeed, with appropriate lags (11 d and 6 For 25 and 29 °C-reared flies), there are high degrees of cross-correlation between the two parameters (Figure 6F & G). To gain further insight into the phenomenon, it would be important to examine mutants with intrinsically altered neuronal excitability and lifespan. Notably, both hyper-excitable *Shaker* (*Sh*) and hypo-excitable *nap*^*ts*^ (*mle*^*napts*^) mutants display shortened lifespans (***Trout and Kaplan, 1970***; ***Reenan and Rogina***, 2008) along with drastically altered ECS discharges (***Lee and Wu, 2006***), flight pattern generation (***Iyengar and Wu (2014)***, unpublished observations), habituation (***Engel and Wu, 1998***).

### Aging-Resilient and -Vulnerable Aspects of Motor Circuit Function in *Drosophila*

The distinct trajectories of aging manifested in the functioning of the various motor circuits described in this study prompted us to ask how high-temperature rearing and oxidative stress, imposed by external environment versus internal milieu, differ in their effects on functional manifestation and aging progression in different motor circuits. Our study clearly delineates the different profiles of the consequences in various physiological functions imposed by oxidative and temperature stressors.

Ambient temperature plays a key role in determining lifespan in ***Drosophila (Loeb and Northrop, 1917***) and other ectotherms (***Van Voorhies and Ward, 1999; Stark et al., 2018***). Under high temperature 29 °C-rearing, lifespan is ∼40-50 % shorter (compared to 25 °C-rearing, median lifespan: 50 vs. 30 d respectively, Figure 1B), and we sought to determine how accurately the age-progression of motor circuit properties could be considered as a simple “compressed” manifestation at a higher temperature. Our data indicate that this notion is largely true for most of the parameters examined here. Parameters that were age-resilient in 25 °C-reared flies were also robustly maintained in 29 °C-reared individuals (SLR and LLR latency and threshold, flight WBF, Figure 7B lighter shade). Similarly, for the two rearing temperatures, even age-vulnerable parameters generally displayed consistent trends in age-dependent changes when scaled for % mortality. This is true particularly for increased DLM firing frequency during flight, increased LLR habituation rate, and decreased ECS seizure threshold (Figure 7B-8, -9, -10 darker shade). Remarkably, bell-shaped age profile of ID spiking was retained in the 29 °C-reared WT population (Figure 7B-11). For the remaining parameters, SLR and LLR refractory periods and DD spiking, 29 °C-rearing appeared to intensify their age-dependence, leading to considerably steeper trends (Figure 7B-3, -6, -12). Our findings suggest that in the shortened lifespans of temperature-stressed individuals, the temporal characteristics of neurophysiological changes are largely retained, albeit “accelerated” according to the degree of lifespan compression. Importantly, these observations give justification to the practice of 29 °C-rearing, a common practice to expedite experiments in *Drosophila* aging studies.

Our findings from *Sod* mutants indicate that oxidative stress exerts strong influences differentially on some of the physiological parameters and the outcomes are distinct from the consequences of high-temperature rearing. Oxidative stress, resulting from inefficient clearance of metabolic reactive oxygen species (ROS, e.g. superoxide anion) and other free radical species, is widely thought as a major contributor to aging processes (***Harman, 1956***, ***1981; Finkel and Holbrook, 2000***). A key class of ROS scavenging enzymes is the Superoxide dismutases which convert superoxide anions into hydrogen peroxide (***Hart et al., 1999; Miller, 2012***). In *Drosophila*, three Superoxide dismutase (Sod) enzymes have been identified: intracellular Cu^2+^/Zn^2+^ Sod1 (encoded by *Sod*, ***Phillips et al. ((1989))***, mitochondrial Mn^2+^ Sod2 (encoded by *Sod2*, ***Duttaroy et al. ((2003)*** also known as *bewildered*, ***Celotto et al. ((2012))***, or extracellular Sod3 (encoded by *Sod3*, ***Jung et al. ((2011))***. Stress resistance and longevity of the fly are greatly compromised when any of the three are disrupted. Indeed, the *Sod* allele described in this study has no detectable enzymatic activity (***Phillips et al., 1989***), and exhibits locomotor defects as well as sensitivity to mechanical-, paraquat- and heat-stress (***Ruan, 2008***).

It is remarkable that *Sod* flies are able to maintain the functionality of the GF pathway throughout their lifespan, as shown by the parameters in the aging-resilient category (SLR and LLR latency and threshold, flight WBF, Figure 7B lighter shade). For several aging-vulnerable properties, *Sod* flies still follow the general profile of age progression with some differences in the amplitude of expression. These are apparent in DLM firing during flight and ID spiking (Figure 7B-8, -11), and more notably, in both accelerated rate of habituation and reduced ECS seizure threshold, closely resembling the relative age-trajectories in WT counterparts (Figure 77B-9 and -10). Nevertheless, striking alterations in *Sod* are seen in the activity-dependent properties of the GF pathway, as reflected in the trajectories of refractory period for both SLR and LLR that are clearly distinct from high-temperature effects (Figure 7B-3, -6). This points to potentially weak links that are sensitive to oxidative stress in the GF-PSI-DLMn circuit for SLR and in the GF afferent processing for LLR.

One conspicuous *Sod* mutational effect revealed in this study is the unexpected, particularly poor performance of the very young age group (< 5% mortality, Figure 7B). These young *Sod* flies displayed remarkably worse flight performance (Figure 11C), longer SLR latency (Figure 2C), greater SLR and LLR RP (Figure 3), increased ECS threshold (Figure 5B), and decreased spiking during seizure discharges (Figures 5C & 6B). Furthermore, a small fraction of aged *Sod* flies (95% mortality) paradoxically perform rather well, comparable to their aged WT counterparts (Figure 7B). A likely explanation for this phenomenon could be the highly stochastic nature of oxidative damages in neural circuits such that a portion *Sod* flies carry variable in-born defects distinct from aging processes. The mutant population undergoes a progressive selection over time in favor of healthier individuals while those with poor circuit performance die off earlier. Consistent with this view, *Sod* mutants reportedly have difficulty eclosing from their pupal case (***Sahin et al., 2017; Woods, 2017***) and the *Sod* lifespan curve lacks the earlier plateau phase (Figure 1B) resulting from much higher mortality in young individuals compared to WT populations.

### Physiological Hallmarks of Aging Progression in Motor Circuits: Quantitative Biomarkers

Identifying biomarkers of aging processes has been an area of active investigation for decades (***Baker and Sprott, 1988; Arking and Wells, 1990; Zou et al., 2000; Butler et al., 2004; Levine et al., 2018***). In *Drosophila*, it is desirable to search for novel physiological aging indices to completement existing cellular and molecular biomarkers of aging, such erroneous expression of the transcription factor genes wingless and engrailed (***Rogina et al., 1997***), accumulation of glycation end-products (***Oudes et al., 1998; Baynes, 2001; Jacobson et al., 2010***), and heat-shock protein (***Landis et al., 2004; Tower, 2011***), as well as modified functional states of the insulin and TOR signaling pathways components (***Kapahi et al., 2004; Partridge et al., 2011***).

Despite the clear differences in motor circuit performance among the different categories of flies examined, two protocols, LLR habituation and electroconvulsively induced seizures, yielded parameters that displayed remarkably consistent age-dependent progressions in normal, high temperature-reared and ROS-stressed populations. Specifically, both acceleration of the habituation rate (Figures 4) and reduction of ECS seizure threshold (Figures 5B) displayed a relatively universal monotonic age-trajectories (Figure 7B-9 and -10). Future studies will further evaluate their suitability as quantitative indices of aging in WT and mutant flies under different environmental conditions or experimental manipulations.

## Methods and Materials

### Fly Stock Maintenance and Lifespan Assays

The Canton-S wild-type (WT) strain and *Sod* mutant allele *Sod*^*1*^ (also known as *Sod*^*n108*^ kept in a balanced stock *Sod*^*1*^ *red/ TM3, Sb Ser*) were maintained on standard cornmeal medium (***Frankel*** *and Brousseau, 1968*). Among *Sod* alleles previously studied (***Campbell et al., 1986; Ruan, 2008***), *Sod*^*1*^ was selected for this study because this allele generated sufficient numbers of flies for crosssectional comparisons across the lifespan. For collecting rare homozygous offspring of *Sod* flies, the stock was grown in half-pint bottles to encourage reproduction. Newly eclosed adults were collected under CO_2_ anesthesia (WT every 1-2 days, *Sod* flies every 2 days) to assemble sufficient sample sizes to initiate lifespan assays. Flies were distributed into food vials (10-14 flies per vial for WT, 7-10 flies for *Sod*) and were transferred to fresh vials every 2 days. For temperature assays, flies were maintained in corresponding 25 °C and 29 °C incubators or at the ambient room temperature of 23 °C. All vials were pre-warmed to the corresponding temperature before fly transfers. Foam plugs were used and the vials were kept horizontally to avoid weaker flies being accidentally stuck to food or cotton plugs. Survivors were counted and dead flies were removed daily. When reported as a function of % mortality, flies were sampled within one day of the age corresponding with % mortality (see Results and Figure 1A).

### Flight Assessment and Indirect Flight Muscle Recording

Preparation, stimulation, recording, and analysis of dorsal longitudinal muscle (DLM) responses have been described previously (***Engel and Wu, 1992; Lee and Wu, 2002; Iyengar and Wu, 2014***). Flies were anesthetized briefly on ice or by light ether exposure and glued to a metal wire between the head and thorax using a nitrocellulose or cyanoacrylate based adhesive (***Gorczyca and Hall, 1984***). Mounted flies were allowed at least 30 min rest in a humid chamber before recording. All electrophysiological and behavioral experiments were carried out at room temperature (21 – 23 °C).

Flights were triggered via a gentle 500 ms air puff generated by an aquarium air pump (Whisper 10-30 Tetra, Blacksburg, VA, USA) pumping air through a 4 mm diameter tube placed approximately 4 mm away from the fly. A 3-way solenoid valve (ASCO scientific, P/N AL4312, Florham Park, NJ, USA) driven by a USB 6210 DAQ card (National Instruments, Austin TX) controlled air flow. Wing sounds were picked up by a high-gain microphone placed approximately 4 mm below the fly. Acoustic signals were digitized by a PC sound card controlled by a LabVIEW 8.6 script. The wing beat frequency was determined using the short-time FFT procedure described in ***Iyengar and Wu (2014)***.

Action potentials from the top-most dorsal longitudinal muscle (#45a, ***Miller (1950))*** were recorded by an electrolytically sharpened tungsten electrode penetrating the flight muscle with a similarly constructed electrode inserted in the abdomen for reference. Signals were accessed with an AC amplifier (Microelectrode AC Amplifier, Model 1800, A-M Systems, Inc., Carlsborg, WA; filter band-width from 10 Hz to 20 kHz). Amplified signals were recorded with pulse code modulation (Neuro-Corder DR-484, Cygnus Technology, Inc., Delaware Water Gap, PA) on videotape at a sampling rate of 44 kHz. Digitization of spiking traces (Figures 1, 6 and Supplemental Figure 2) was carried out by a digital acquisition card (USB 6210) controlled by custom-written LabVIEW scripts.

### Giant Fiber Pathway Stimulation

As previously described (***Engel and Wu, 1996***), a pair of uninsulated tungsten electrodes were inserted in the eyes (anode in left eye) to pass stimulation of 0.1 ms duration (Isolated Pulse Stimulator, Model 2100, A-M Systems, Inc., Carlsborg, WA). Long-latency and short-latency response (LLR and SLR) thresholds were first determined using increasing stimulation intensity with an interstimulus interval of 30 s. For the experiments involving LLR (twin-pulse refractory period and habituation), stimulus intensity was set at least 0.5 V below the SLR threshold. For SLR-related trials, stimulus intensity was set at 20 V, which was well above the SLR thresholds in all flies tested (see ***Engel and Wu (1992)*** for details). To determine the refractory period of LLR and SLR, twin-pulse stimuli were given every 10 s, with the stimulus interval decreasing stepwise (step size: 5 - 10 ms for LLR, 0.5 - 1 ms for SLR) from a starting interval (100 - 800 ms for LLR, 10 - 20 ms for SLR). When a given inter-stimulus interval failed to trigger a second DLM spike in one or two trials, a 3 - 5 min rest was allowed before a train of six twin-pulses of the same inter-stimulus interval were delivered at 0.1 Hz between twin pulses. If all six trials failed, then the previous stimulus interval was recorded as the refractory period. The ability of GF to follow high-frequency stimulation (resp. 30 p.) was determined as described previously (***Gorczyca and Hall, 1984***; ***Engel and Wu, 1992***). Three trains of 10 pulses at 200 Hz with 10 s in between were delivered and the number of responses was counted.

### Habituation of GF-Mediated LLRs and Electroconvlusive Stimulation-Evoked Seizures

For testing habituation rate, two trials of 100-pulse LLR stimulation trains at a given frequency (1, 2, and 5 Hz) were delivered with a 10 min interval between trials. The criterion for reaching habituation was the occurrence of the first five consecutive LLR failures (F5F, ***Engel and Wu (1996))***. The larger F5F value of the two trials was recorded for the fly tested. Upon reaching habituation criterion, an air puff dishabituation stimulus was delivered to confirm habituation (see ***Engel and Wu (1996)*** for additional details).

High-frequency electroconvulsive stimulation (0.1-msec pulses at 200 Hz for 2 s at a particular voltage 15 - 80 V) were delivered across the brain to induce a stereotypic electroconvulsive seizure (ECS) discharge pattern (see ***Lee and Wu (2002)*** for details). After high-frequency stimulation, test pulses of 20 volts or higher were sometimes delivered at 1 Hz to examine the failure and recovery of the GF pathway motor response. An interval of at least 10 min was allowed between each ECS to avoid the effect of refractoriness (***Lee and Wu, 2002***).

For follow-up experiments of DLM spike patterning during ECS discharges (Figure 6 and Supplemental Figure 2), spikes were detected and their timing recorded using custom-written Matlab scripts (***Iyengar and Wu, 2014; Lee et al., 2019***). Firing rate was measured by the instantaneous firing frequency (ISI^−1^, defined as the reciprocal of the inter-spike interval, ISI between successive spikes), and firing regularity was quantified by the instantaneous coefficient of variation (CV_2_ see ***Holt et al. ((1996)***; ***Lee et al. ((2019))***. Poincaré trajectories were constructed by plotting the ISI^−1^ of a spike interval against the ISI^−1^ of the subsequent interval. Averaged ISI^−1^ vs CV_2_ plots were constructed as described in ***Lee et al. ((2019)***.

The neurophysiological properties of each fly were tested in the following sequence with 5 min rest in-between to minimize interference between protocols: flight, LLR and SLR thresholds, LLR refractory period, LLR habituation, SLR refractory period, ability of SLR to follow high frequency stimulation (30 pulses at 200 Hz), and ECS-evoked seizure discharges. Due to the weakness of aged flies and in some cases the absence of LLRs in the last 5% survivors, not all of the flies went through every protocol. If the LLR was absent, all SLR-related protocols were still examined. Generally, all electrophysiological protocols took less than one hour to complete for a single fly.

### Statistical Analysis

For each age group tested (1, 5, 50 and 95% mortality), between 7 and 12 flies were generally tested. We have found that this sample size range is often sufficient to draw initial conclusions, based on our previous experiences studying ion channel and 2nd messenger system mutants (***Engel and Wu, 1992, 1996; Lee and Wu, 2006***; ***Iyengar and Wu, 2014***). Individual flies were considered biological replicates, and all recorded data points were included in the analysis. Due to the non-normal distribution of datasets, the non-parametric Kruskal-Wallis ANOVA test was used to determine crosssectional differences between physiological parameters of flies from respective genotypes. The rank-sum test (with Bonferroni correction applied) was used as a post hoc test. Age-dependent trends in physiological parameters were identified by computing the non-parametric Spearman’s rank correlation coefficient (r_s_) (***Sokal and Rohlf, 1969***). In the source data files accompanying each figure, the sample mean, median, standard deviation, coefficient of variation and the 25 – 75 %tile interval are listed. To assess the robustness of the respective statistics, a bootstrap resampling approach (Efron, 1981) with 1000 replicates was used to estimate the 95%tile confidence intervals. All statistical analyses were performed in Matlab (r2019b).

## Acknowledgments

We thank Xiaomin Xing, Scott Woods, Tristan O’Harrow and Atsushi Ueda for critical discussions during the projects, Jeff Nirschl, Jordan Imoehl, Kelle Goranson, and Amy Hoehne for assistance in data collection. This research was supported by a University of Iowa Biological Sciences Funding Program Grant and NIH grants, GM088804 AG047612, and AG051513 to CFW; and by an NIH NRSA Fellowship NS082001 to AI

## Supplemental Information

**Figure S1.**
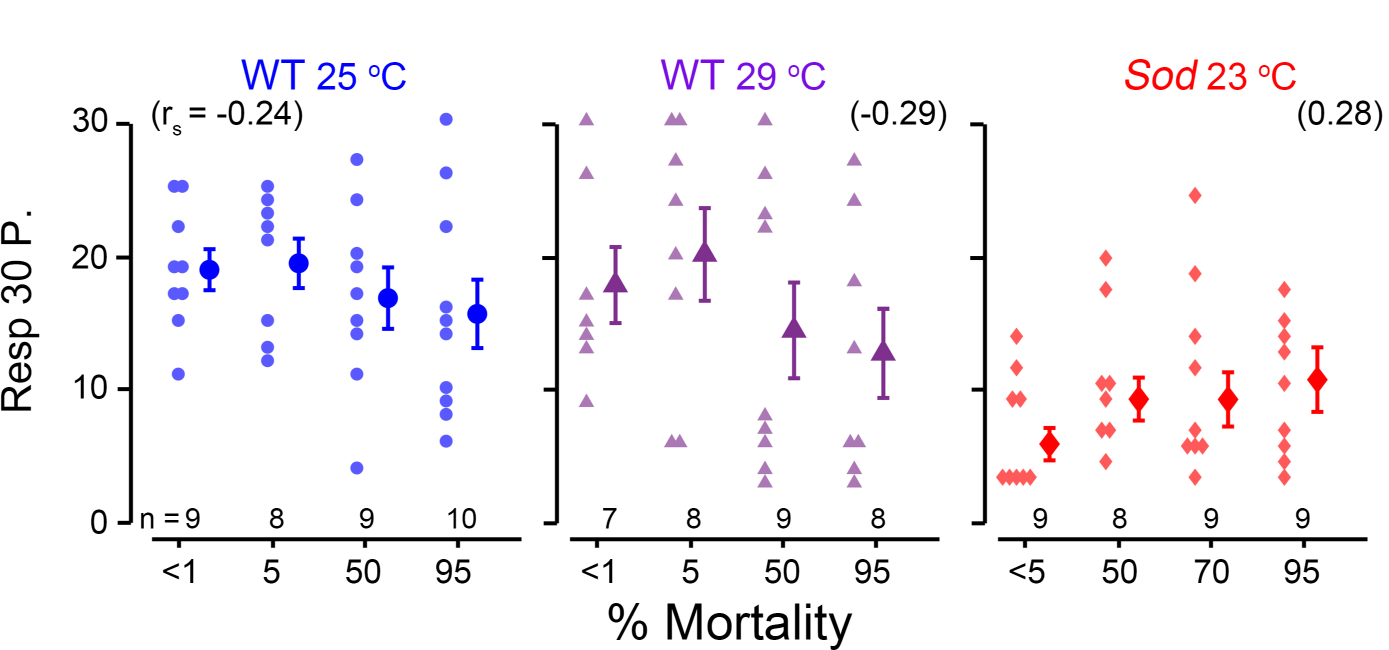
Giant-fiber SLR properties across the lifespan: resp. 30 p. protocol. The resp. 30 p. protocol measures the ability of the GF pathway to follow high-frequency stimulation. Three trains 10 stimuli are delivered at 200 Hz with a 10 s interval between trains. The number of responses was recorded, with a higher response rate corresponding to a better ability to follow high-frequency stimulation. Sample sizes as indicated for each age group, and the Spearman rank correlation (r_s_) is shown above. No population displayed statistical significance age-dependent trend (i.e. p > 0.05).

**Figure S2.**
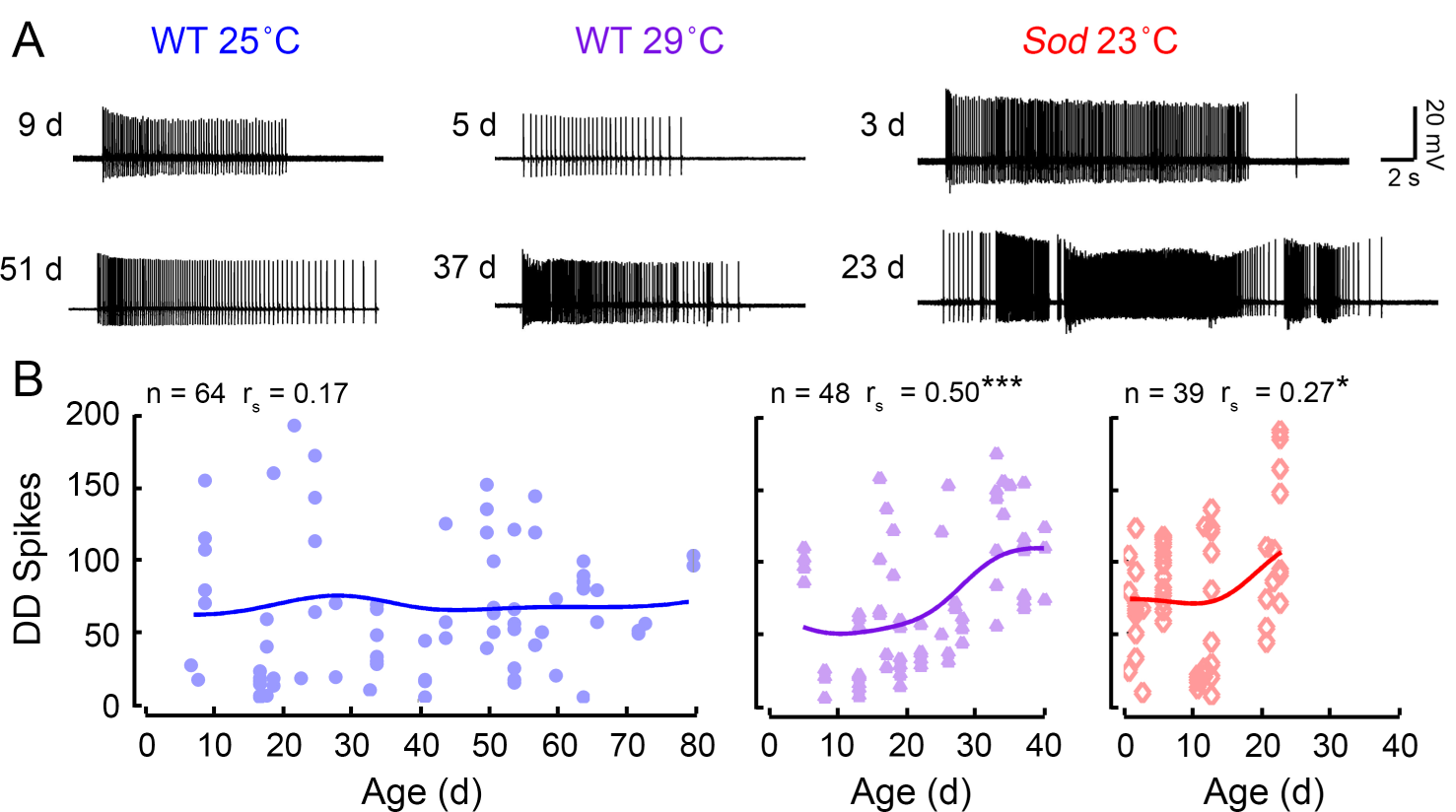
Spiking activity of delayed seizure discharges across the lifespan. (A) Example traces of DD spiking in young (top row) and aged (bottom row) flies. (B) Scatterplot of the spike count during the DD. Trend-line indicates the Gaussian kernel running average as computed in Figure 1. The sample size and age-dependent rank correlation (r_s_) are indicated above. (* p < 0.05, *** p < 0.001).

## Notes

### Competing Interest Statement

The authors have declared no competing interest.

### Summary of Updates

Title & Abstract Changed

