## Supplementary figures and images for "Distinct aging-vulnerable trajectories of motor circuit functions in oxidation- and temperature-stressed *Drosophila*"

### Supplemental Figure 1

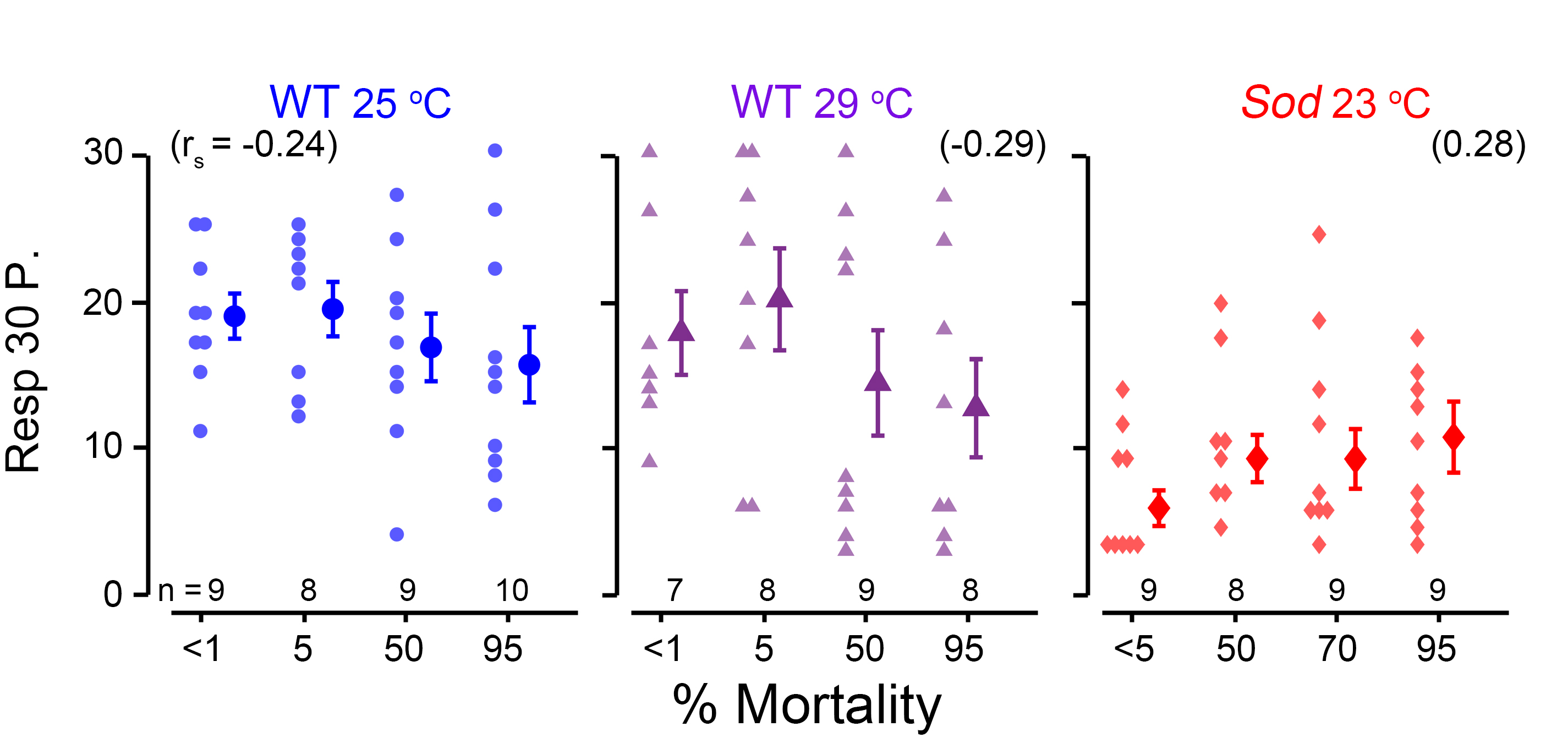

### Supplemental Figure 2

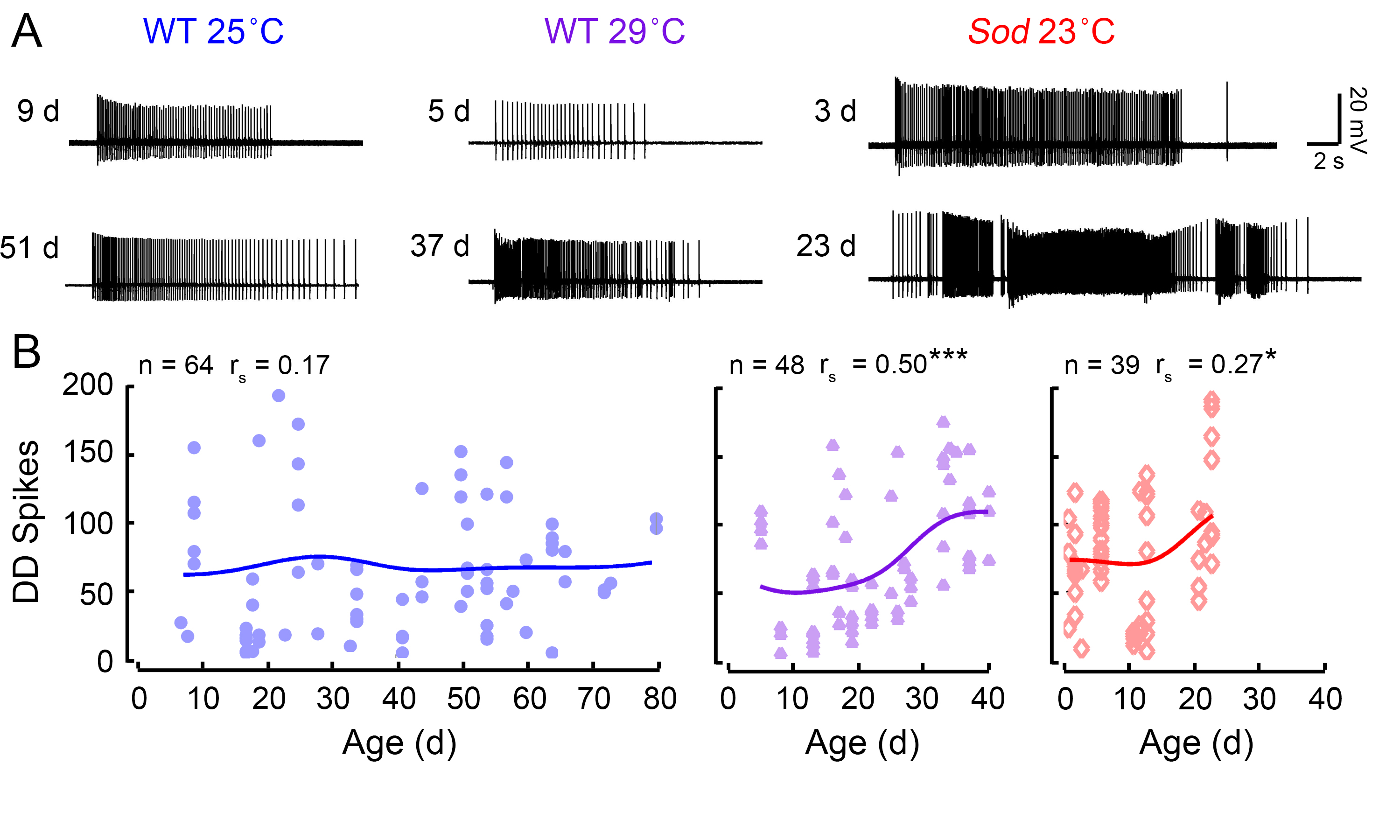
